# Low-Background Cancer Imaging With a Bioorthogonal Fluorescence Probe and Engineered Reporter Enzyme Bearing a Targeting Moiety

**DOI:** 10.1101/2025.03.04.641560

**Authors:** Ziyi Wang, Ryosuke Kojima, Rikuki Kiji, Kyohhei Fujita, Ryo Tachibana, Taku Uchiyama, Yoshihiro Minagawa, Tadahaya Mizuno, Kiyohiko Igarashi, Hiroyuki Noji, Mako Kamiya, Yasuteru Urano

## Abstract

Combinatorial use of an antibody-reporter enzyme conjugate and a fluorescence probe activated by the enzyme is a powerful strategy for fluorescence-guided cancer surgery. However, conventional probes for typical reporter enzymes are insufficiently bioorthogonal, resulting in high background signals in non-target tissues. We screened a library of HMRef (rhodol derivative)-based fluorescence probes bearing various sugar moieties, and discovered that HMRef-β-D-fucose is bioorthogonal in mammalian systems, but is activated by a metagenomic glycosidase, Td2F2. Directed evolution generated a mutant with a *k*_cat_/*K*_m_ value for HMRef-β-D-fucose of 3.3 x 10^5^/M/sec, 7.3 times higher than that of wild-type Td2F2 and comparable to that of β-galactosidase (LacZ) with a corresponding probe. Theoretical calculation suggested that E296G mutation in Td2F2 causes structural changes that facilitate the probe’s access to the enzyme’s active site. In a proof-of-concept study, cancer cells were visualized with a minimal background in the mesentery of a mouse model of peritoneally disseminated human-ovarian-cancer-derived SKOV-3 cells, which endogenously express HER2, by using HMRef-β-D-fucose together with engineered Td2F2 conjugated/fused to a HER2-binding antibody/nanobody.

## Introduction

Complete dissection of cancer during surgery is critical to prevent cancer recurrence, and fluorescence-guided surgery is one of the most promising approaches to assist in achieving this^1–6^. For example, fluorogenic probes targeting cancer-specific hydrolases (e.g. peptidases such as γ-glutamyltransferase^7^ and dipeptidyl peptidase-IV^8^, and glycosidases such as α-mannosidase 2C1^9^) are powerful tools to achieve cancer imaging with a high signal-to-noise ratio by leveraging enzymatic turnover to amplify the signal. However, it is not always possible to find an appropriate cancer-specific enzymatic activity, depending on the cancer cell type to be visualized.

A powerful strategy to overcome these limitations is to deliver a reporter enzyme selectively to cancer cells and then administer a fluorogenic probe that is a substrate of the enzyme^10,11^ (Figure 1a). This method is conceptually similar to directed enzyme prodrug therapy (DEPT)^12,13^, which has been extensively explored for targeted drug activation. A typical method to deliver the reporter enzyme is to conjugate it with an antigen-binding protein targeting a cancer-specific cell-surface antigen. For example, a conjugate of β-galactosidase (expressed from *lacZ*) with avidin, which binds to lectins displayed on cancer cells, has been used to target cancer cells expressing high levels of lectins, and subsequent administration of a fluorogenic probe that is a substrate of this enzyme enabled clear visualization of the cancer cells^14^. This kind of approach can target a broad range of cancer cells, because a variety of cancer-specific cell-surface antigens can be harnessed^15,16^. A key advantage of this approach compared to the use of antigen-binding proteins directly labeled with fluorophores is that large amounts of fluorophore are generated per antigen as a result of enzymatic turnover (as is the case when using fluorescence probes against endogenous hydrolases), leading to bright imaging of cancer cells.

**Figure 1.**
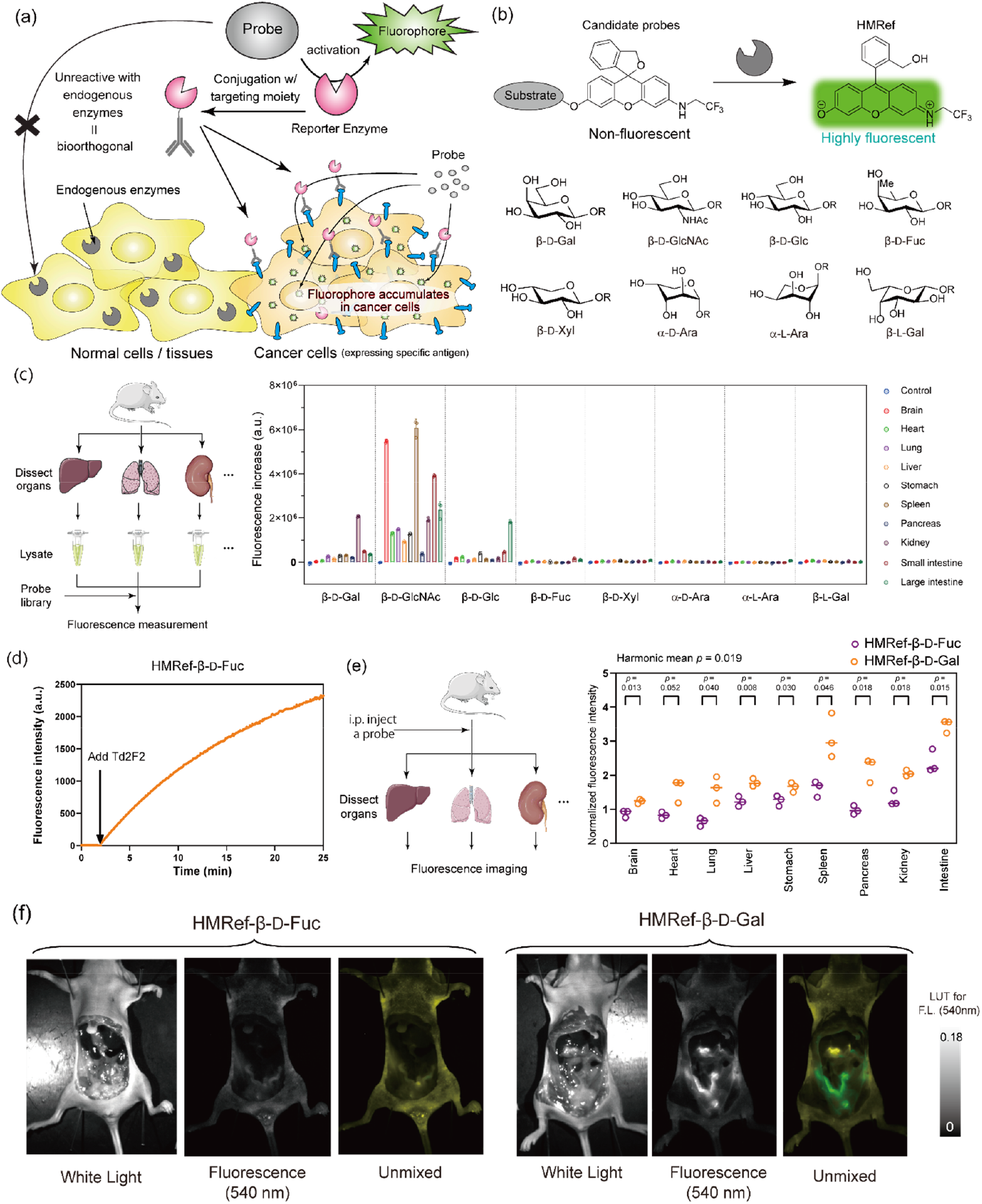
Schematic illustration of bioorthogonal fluorescence probe and reporter enzyme. (**a**) Principle of the cancer imaging technique presented in this study. Substrates of typical reporter enzymes are often activated by endogenous enzymes with similar activities. Therefore, we aimed to find a bioorthogonal fluorescence probe that is not activated by endogenous enzymes and a non-endogenous reporter enzyme that hydrolyzes it, thereby enabling selective cancer imaging. (**b**) Fluorescent probe library used in this study. We tested the bioorthogonality of a series of HMRef derivatives bearing different sugar moieties (β-D-galactose (Gal), β-D-N-acetylglucosamine (GlcNAc), β-D-glucose (Glc), β-D-fucose (Fuc), β-D-Xylose (Xyl), α-D-arabinose (Ara), α-L-Ara, and β-L-Gal in order to identify probes bearing a sugar moiety that is not hydrolyzed by mammalian endogenous enzymes but can be hydrolyzed by enzymes from other species. (**c**) Work flow and the results of the screening with mouse organ lysate (Jcl:ICR). Fluorescence increases at 60 min after the addition of fluorescence probes to organ lysates in PBS (pH 7.4) [probe] = 1 μM, [lysate] = 0.5 mg protein/mL. (**d**) Time course of fluorescence intensity of HMRef-β-D-Fuc in the presence of Td2F2. [probe] = 1 μM, [Td2F2] = 100 nM. (**e**) Work flow and the results of *in vivo* evaluation of HMRef-β-D-Fuc and HMRef-β-D-Gal after i.p. injection of the probes. BALB/c-nu/nu mice were i.p. injected with 150 μL of 1 mM probes, and the main organs were dissected after 30 min for imaging (Ex/Em = 490 nm/515 nm-long pass). (Note that the fluorescence includes the autofluorescence of each organ.) *P*-values: two-tailed Welch’s t-test (n = 3). The harmonic mean *p*-value^43^ reflecting the multiple comparison is also shown. The original imaging data can be found in Supplementary Figure 2. (**f**) Fluorescence imaging of the peritoneal cavity of Balb-c nu/nu mice i.p. injected with 150 µL of 1 mM HMRef-β-D-Fuc or HMRef-β-D-Gal. After 1 hour, the mice were sacrificed and subjected to imaging after dissecting out the intestines. Ex/Em = 490 nm/515 nm-long pass.

However, probes used with typical reporter enzymes are often insufficiently bioorthogonal, resulting in high background signals due to activation by endogenous enzymes. For example, β-galactosidase is expressed in various tissues in mammals (encoded by GLB1 in humans) and indeed is so highly expressed in senescent cells that β-galactosidase probes are used for visualizing them^17,18^. Probes used with other reporter enzymes such as alkaline phosphatase^19^, nitroreductases^20,21^, esterases^22^, and horseradish peroxidase^23^ are also insufficiently bioorthogonal.

Here, to address this issue, we screened our glycosidase-targeting fluorescence probe library^9^, and discovered a probe bearing β-D-fucose (HMRef-β-D-fucose) that is resistant to endogenous mammalian enzymes. Moreover, we found that this probe is activated by a metagenomic enzyme Td2F2^24^ belonging to the GH1 family. We employed fluorescence activated cell sorting (FACS)-based high-throughput directed evolution to obtain a Td2F2 mutant with dramatically increased activity, and confirmed that conjugates of this engineered enzyme with various cancer-antigen-binding proteins, in combination with HMRef-β-D-fucose, can clearly visualize live cancer cells *in vitro*. Furthermore, as a proof-of-concept, we showed that cancer cells can be clearly visualized with a minimal background in the mesentery of a mouse model of peritoneally disseminated human-ovarian-cancer-derived SKOV-3 cells using the combination of HMRef-β-D-fucose with engineered Td2F2 chemically conjugated/genetically fused to Trastuzumab/anti-HER2 nanobody as the targeting moiety.

### Screening to discover a bioorthogonal probe and reporter enzyme combination

In order to achieve cancer imaging with a reduced background, it is necessary to develop fluorescence probes that are inert to enzymes contained in normal tissues (Figure 1a). Therefore, we screened our previously reported HMRef^25^ (rhodol derivative)-based fluorescence probe library bearing different sugar moieties^9^ (Figure 1b). Conjugation of a sugar to the oxygen atom of HMRef essentially nullifies the absorption and fluorescence of HMRef in the visible region due to intramolecular spirocyclization (leading to π-deconjugation of the xanthene moiety), and cleavage of the sugar leads to the release of HMRef, which shows bright green fluorescence^25^ (Figure 1b). Since the number of sugars that can be digested by the human/mammalian body is limited, we hypothesized that we would be able to identify probes bearing a sugar moiety that is not hydrolyzed by mammalian endogenous enzymes, but can be hydrolyzed by enzymes from other species.

We tested the reactivity of the 8 probes shown in Figure 1b by mixing them with mouse organ lysates (Figure 1c). We found that conjugates of HMRef with β-L-galactose (Gal), β-D-xylose (Xyl), α-D-arabinose (Ara), α-L-Ara, and β-D-fucose (Fuc) did not show a significant fluorescence increase after co-incubation for 60 minutes with most of the tissue lysates, whereas HMRef conjugated with β-D-Gal, β-D-glucose (Glc), and β-D-N-acetylglucosamine (GlcNAc) were hydrolyzed by the organ lysates, leading to a higher fluorescence increase. The latter finding is consistent with the fact that the probe HMRef-β-D-Gal, used as a substrate of one of the most commonly used reporter enzymes, β-galactosidase^26^, exhibits background activation in various tissues.

Next, we set out to find a glycosidase that can hydrolyze the candidate bioorthogonal probes. We focused on a metagenomic glycosidase Td2F2^24^ belonging to the GH1 family as a candidate reporter enzyme for the β-D-Fuc-bearing substrate for the following reasons. (1) It can be easily expressed and purified with standard methods. (2) A relatively high *k*_cat_ at physiological conditions has been experimentally established. (3) It is monomeric, simplifying bioconjugation. (4) The crystal structure has been determined^27^, which would be helpful for engineering the enzyme. Gratifyingly, when we tested the reaction of HMRef-β-D-Fuc with recombinant Td2F2 expressed in *E. coli* (Supplementary Figure 1), a clear fluorescence increase was indeed observed (Figure 1d).

Motivated by this finding, we further assessed the bioorthogonality of HMRef-β-D-Fuc in comparison with HMRef-β-D-Gal, considering the wide usage of β-galactosidase as a reporter enzyme. In a systemic comparison, HMRef-β-D-Fuc showed markedly lower fluorescence than HMRef-β-D-Gal in various organs after i.p. injection (Figure 1e, Supplementary Figure 2). Furthermore, when the peritoneal cavity was directly visualized after i.p. injection of the probes, HMRef-β-D-Fuc showed almost no fluorescence, while significant fluorescence was observed with HMRef-β-D-Gal (Figure 1f). We also confirmed that HMRef-β-D-Fuc did not react with human serum (Supplementary Figure 3). Taking also into account that this probe seldom reacts with human-derived cell lines^9^, we decided to evaluate the potential of the HMRef-β-D-Fuc and Td2F2 pair for cancer imaging, in comparison with HMRef-β-D-Gal and β-galactosidase.

### Cancer imaging using wild-type (WT) Td2F2/HMRef-β-D-Fuc

To evaluate the cancer-imaging ability of the selected enzyme-probe pair, we conjugated the enzyme to proteins capable of binding to cancer-specific antigens and used the conjugates to target cultured cancer cells. Specifically, we conjugated Td2F2 to avidin, which binds to lectins expressed on cancer cells^14,28,29^, or Trastuzumab, which is an antibody recognizing HER2^30^ (expressed on various cancer cells including breast cancer and ovarian cancer^31,32^) (Figure 2a). For conjugation with avidin, we biotinylated Td2F2 with biotin-succinimidyl ester (SE) and mixed it with avidin in an approximately 1:1 ratio (one Td2F2 molecule per one avidin tetramer). For conjugation with Trastuzumab, we artificially installed cysteine (Cys) at the N-terminus of Td2F2 for labeling, since there is no surface-exposed Cys in Td2F2 (Supplementary Figure 4), and conjugated this with Trastuzumab via a 3-sulfo-N-succinimidyl 4-(N-maleimidomethyl)cyclohexane-1-carboxylate (Sulfo-SMCC) linker. For comparison, we also prepared avidin-β-galactosidase (expressed from *lacZ*) and Trastuzumab-β-galactosidase conjugates in a similar manner. It should be noted that β-galactosidase forms a tetramer with multiple surface-exposed Cys residues (Supplementary Figure 4). In contrast, the monomericity and the site-specific installation of Cys for labeling of Td2F2 should be beneficial for bioconjugation.

**Figure 2.**
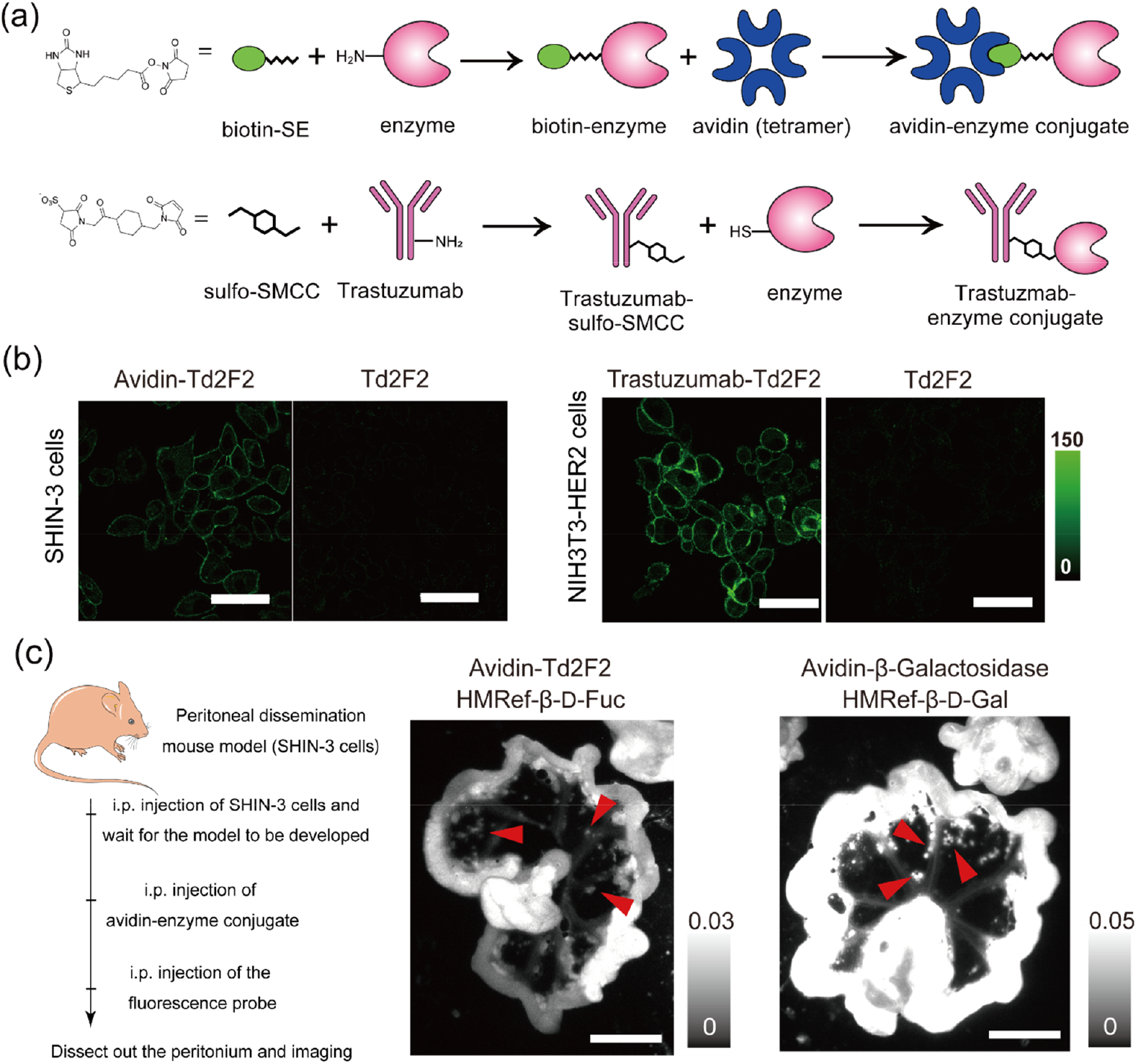
Cancer imaging with HMRef-β-D-Fuc and Td2F2 WT, and comparison with HMRef-β-D-Gal and β-galactosidase (expressed from *lacZ*). **(a)** Schematic illustration of the construction of avidin-enzyme conjugate and Trastuzumab-enzyme conjugate, exemplified by the Td2F2 conjugate (note that β-galactosidase forms a tetramer). For the sulfo-SMCC-based conjugation of Td2F2, a cysteine residue was artificially installed at the N-terminus of Td2F2 because there is no available Cys on the surface of the Td2F2 molecule (Supplementary Figure 4). **(b)** Live cell imaging of SHIN-3 cells (galectin-positive) and NIH3T3-HER2 cells with avidin-Td2F2 and trasutuzumab-Td2F2, respectively. Cells were treated with 100 nM Td2F2 conjugated with the corresponding antigen-binding protein or with Td2F2 only for 16 hours. After washing, 1 µM HMRef-β-D-Fuc was applied. Then imaging was conducted after 1 hour with a confocal fluorescence microscope. Ex/Em: 488 nm/498-550 nm. Scale bar: 50 µm. **(c)** Fluorescence imaging of SHIN-3 cells disseminated in mouse peritoneal cavity. 500 μL PBS containing 500 nM avidin-Td2F2 or 100 nM avidin-β-galactosidase was i.p. injected into Balb-c nu/nu bearing peritoneally disseminated SHIN3 cells. Sixteen hours later, 150 μL of 1 mM fluorescence probe (HMRef-β-D-Fuc for avidin-Td2F2, HMRef-β-D-Gal for avidin-β-galactosidase) was injected. One hour later, the mesentery was dissected out and subjected to fluorescence imaging. Ex/Em = 490 nm/515 nm-long pass, Scale bar = 10 mm. The red arrowheads show examples of clumps of cancer cells (other spots like these are also clumps of cancer cells).

We first tested combinations of Td2F2 conjugates and HMRef-β-D-Fuc for live cell imaging. Avidin-Td2F2 was used to image human-ovarian-cancer-derived SHIN-3 cells, which highly express lectins recognized by avidin^14^. Trastuzumab-Td2F2 was applied for imaging human-ovarian-cancer-derived SKOV-3 cells, which endogenously express high levels of HER2. After manual wash out of surplus Td2F2 conjugate, HMRef-β-D-Fuc was applied, and fluorescence appeared in both cases (Figure 2b). Almost no fluorescence was observed when Td2F2 without the antigen-binding moiety was used, because the enzyme was washed out before the probe application. This result confirmed the applicability of these enzyme-probe combinations for target-antigen-dependent cancer cell imaging. Next, we examined *in vivo* cancer imaging. We chose a model of peritoneal dissemination, as this is an advanced symptom of several visceral cancers including ovarian cancer. Specifically, a mouse model with peritoneal dissemination of SHIN-3 cells was prepared and i.p. injected with avidin-enzyme conjugate (Td2F2 or β-galactosidase). After waiting for the surplus conjugates to wash out naturally, the corresponding fluorescence probe (HMRef-β-D-Fuc for Td2F2, and HMRef-β-D-Gal for β-galactosidase) was i.p. injected. Imaging of the dissected mesentery showed that cancer cells were clearly visualized with a low background with the Td2F2/HMRef-β-D-Fuc system (Figure 2c). However, the signal from the cancer cells was low compared to that obtained with the β-galactosidase/HMRef-β-D-Gal system (Figure 2c; note that a 5 times greater amount of the Td2F2 conjugate was used). Therefore, we set out to increase the enzymatic activity of Td2F2 by means of directed evolution.

### Engineering of Td2F2 to enhance its activity

To enable efficient engineering of Td2F2 in a pooled format, we employed high-throughput fluorescence-activated cell sorting (FACS) of *E. coli* cells expressing Td2F2 mutants. A key experimental requirement is one-to-one correspondence between the activity (phenotype) of the mutant and its genotype at the single-cell level, even in a pooled format^33,34^. However, with conventional small-molecule fluorescence probes, including HMRef-β-D-Fuc, the activated fluorophore can leak out of the cells, leading to loss of correspondence between genotype and phenotype (Supplementary Figure 5). To address this point, we synthesized a new probe, SPiDER-β-D-Fuc. This probe, which differs from HMRef-β-D-Fuc in having a fluoromethyl group ortho to the hydroxyl group of rhodol, becomes fluorescent after reaction with Td2F2 and simultaneously generates a highly electrophilic quinone methide that reacts with intracellular proteins^35^. We experimentally confirmed that this leads to intracellular trapping of the fluorescent product, enabling labeling of *E. coli* cells with single-cell resolution and sorting of each mutant by FACS with retention of the correspondence of phenotype and genotype (Figure 3a, Supplementary Figure 5). With a suitable high-throughput assay and isolation procedure in hand, we next utilized error-prone PCR with the full-length Td2F2 as a template to prepare a pool of expression cassettes coding more than 10^7^ random mutants of Td2F2. The variants were expressed in *E. coli* cells, and SPiDER-β-D-Fuc was introduced into the cells aided by the lower osmotic pressure of the surrounding medium^36^. *E. coli* cells showing the highest green fluorescence were isolated by FACS, and the Td2F2 coding region was re-amplified to prepare a new library. During four rounds of this cycle, the fluorescence intensity of the brightest cells (as well as the proportion of fluorescent cells) increased, suggesting that highly active Td2F2 mutants were enriched in the sorted population (Figure 3b). Finally, the evolved Td2F2 mutants were individually expressed from 310 colonies and assayed in separate wells. Gratifyingly, we succeeded in identifying multiple mutants with potent enzymatic activity, including Mutant A (E27G, I34T, F243L, E296G) (Figure 3c), which showed significantly higher *k*_cat_/*K*_m_ values with both HMRef-β-D-Fuc (7.3-fold higher) and SPiDER-β-D-Fuc (8.4-fold higher) compared to WT Td2F2 (Figure 3d, Supplementary Figure 6). Notably, the *K*_m_ values of the probes were greatly reduced, which would be especially beneficial for *in vivo* applications where maintaining a high probe concentration is difficult. The *k*_cat_/*K*_m_ value of Td2F2 Mutant A with HMRef-β-D-Fuc was comparable to that of β-galactosidase expressed from *lacZ* with HMRef-β-D-Gal, suggesting that our enzyme/probe pair would be suitable for cancer imaging applications.

**Figure 3.**
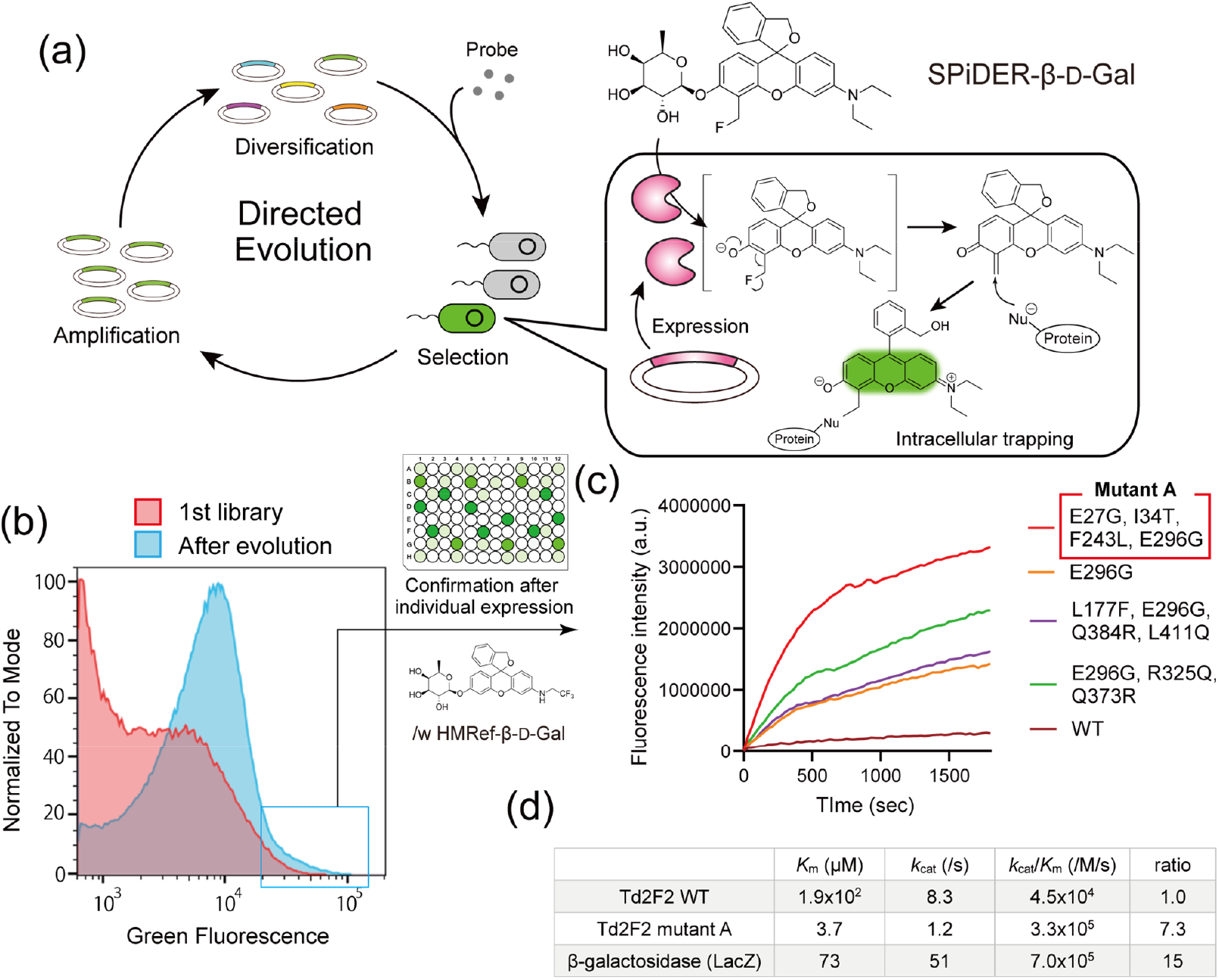
Directed evolution of Td2F2 aimed at increasing the activity for activation of fluorogenic probes bearing β-D-Fuc. (**a**) Schematic illustration of the directed evolution of Td2F2. For this, we developed a new probe, SPiDER-β-D-Fuc, that forms a highly electrophilic quinone methide after hydrolysis of the β-D-Fuc moiety. The methide reacts with intracellular proteins, assuring one-to-one correspondence between the enzymatic activity of each Td2F2 mutant and the fluorescence intensity of the *E. coli* cells with single-cell resolution. (**b**) Monitoring the outcome of directed evolution with FACS. After 4 rounds of concentration of highly active mutants by FACS sorting, *E. coli* cells showing brighter fluorescence than cells expressing WT Td2F2 were observed. (**c**) The assay results of individual mutants in the homogeneous solution. After transforming *E. coli* cells with the plasmid library encoding Td2F2 mutants (recovered from the cells sorted in (**b**)), we picked up 310 colonies and induced protein expression to test the activity of individual mutants in homogeneous solution. Each cell lysate was diluted in 100 mM NaPi buffer (pH 7.4) and reacted with 1 μM HMRef-β-D-Fuc. The fluorescence increase was monitored with a plate reader. The sequences of the most active mutants are shown. (**d**) Enzymatic activity parameters of Td2F2 WT and Mutant A (E27G, I34T, F243L, E296G, see (**c**)). The *k*_cat_/*K*_m_ value of Mutant A towards HMRef-β-D-Fuc is comparable to that of β-galactosidase (from *lacZ*) towards HMRef-β-D-Gal.

Analysis of the sequences of the mutants showing high reaction rates (including Mutant A) suggested that E296G is a critical mutation for increased activity (Figure 3c). Quantum mechanics/molecular mechanics (QM/MM) calculation of the structure of Td2F2 mutants and the probes (utilizing the crystal structure of WT Td2F2 with the corresponding substrate as a template^27^) (Supplementary Data 1-6) suggested that the E296G mutation structurally facilitates the probe’s access to the enzyme’s active site. E296 forms a salt bridge with R325, which is thought to reduce the accessibility of the active site, so the E296G mutation may eliminate this interaction (Supplementary figure 7). The QM/MM calculation also suggested that this mutation contributes to the bond-stabilization energy of the encounter complex of the probes and the enzyme (Supplementary figure 7). This result is consistent with the experimentally confirmed decrease of *K*_m_ of the probes with the mutant, strongly supporting the putative structural basis of the enhanced performance of the mutant.

### Application of the Td2F2 mutant for cancer imaging

To test the performance of Td2F2 Mutant A, we conjugated it to Trastuzumab using the same method described above and applied it for imaging of live NIH3T3-HER2 cells. This mutant imaged HER2-positive cells with a much stronger signal than Td2F2 WT (Figure 4a). Next, we compared the performance of Td2F2 Mutant A, Td2F2 WT, and β-galactosidase, each of which was conjugated to Trastuzumab, in a mouse model with peritoneal dissemination of SKOV-3 cells. As in the case of the avidin conjugate discussed above, the conjugates were i.p. injected into the mice. After waiting for the surplus conjugates to wash out naturally, HMRef-β-D-Fuc (for the Td2F2 system) or HMRef-β-D-Gal (for the β-galactosidase system) was i.p. injected. Fluorescence imaging of the dissected mesentery clearly showed that the combination of HMRef-β-D-Fuc and the Trastuzumab conjugate of Td2F2 Mutant A achieved vivid cancer imaging with a fluorescence intensity comparable to that of the HMRef-β-D-Gal and β-galactosidase conjugate system, while affording a minimal background signal (Figure 4b). The cancer-to-normal-tissue (intestine) ratio in mice injected with the Td2F2 Mutant A system was significantly larger than that in the case of β-galactosidase-treated mice (Figure 4c. Note that the quantification includes tissue autofluorescence, so a ratio of less than 1 does not mean the probes are hydrolyzed more in the intestines, especially for the β-D-Fuc/Td2F2 system. See Supplementary Figure 8 for unmixed images). In the case of the β-galactosidase system, significant background signals from multiple tissues including the intestines and uterus were observed, whereas this was not the case with the Td2F2-based system (Supplementary Figure 8). Since the background signals were observed even in the absence of the enzyme conjugate (Figure 1f), the background should be mainly due to insufficient bioorthogonality of HMRef-β-D-Gal, rather than insufficient wash-out of the conjugate. These results establish the effectiveness of the newly developed engineered Td2F2/HMRef-β-D-Fuc pair for cancer imaging.

**Figure 4.**
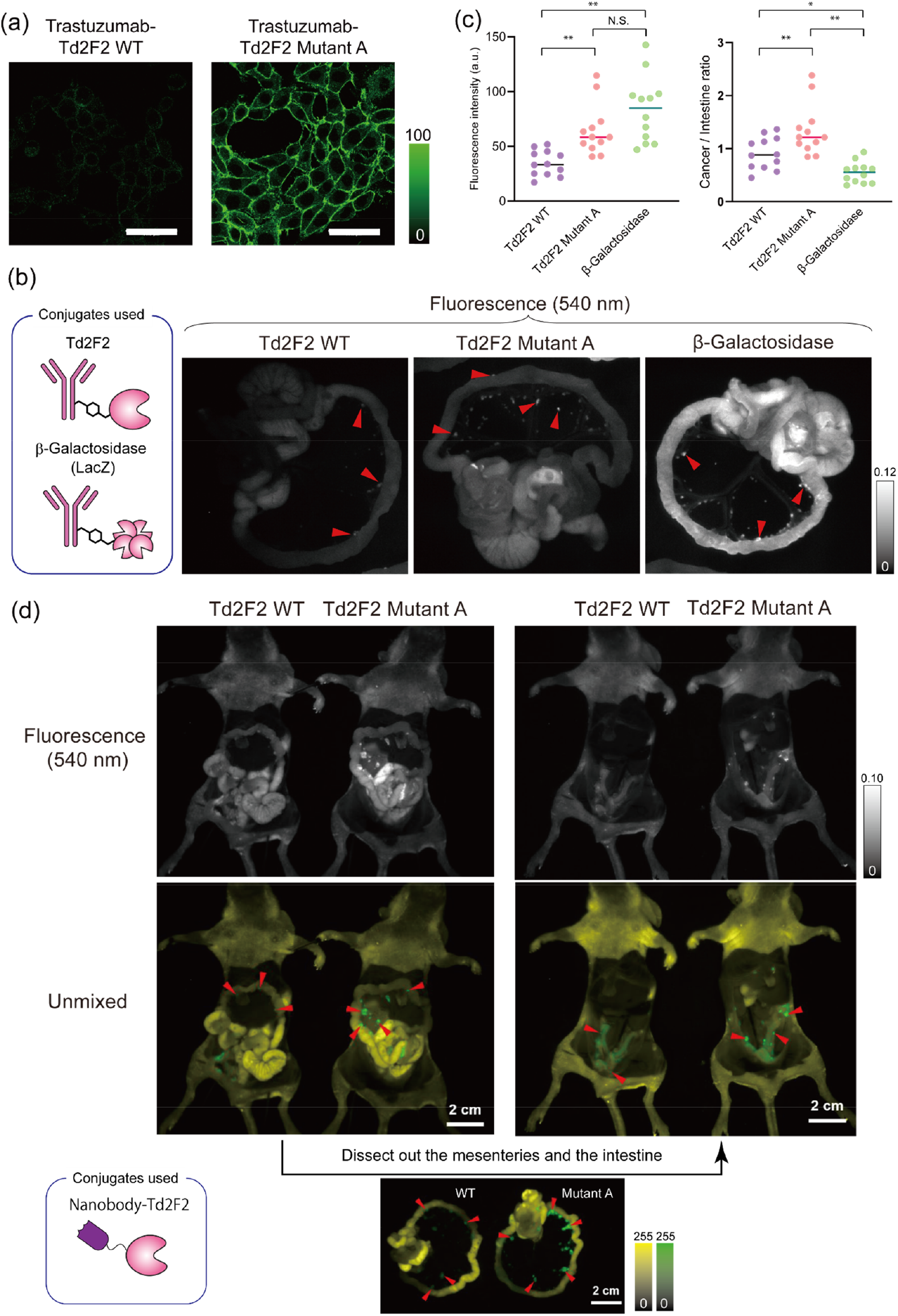
Cancer imaging with HMRef-β-D-Fuc and the conjugates of the Td2F2 (mutant) and antigen-binding proteins. (**a**) Comparison of Td2F2 WT and Td2F2 Mutant A (E29G, I34T, F243L, E296G) for live-cell imaging. NIH3T3-HER2 cells were treated with 100 nM of the conjugates of Td2F2 WT/Mutant A and Trastuzumab for 16 hrs. After washing, 1 µM HMRef-β-D-Fuc was applied, and imaging was conducted 1 hour later with a confocal fluorescence microscope. Ex/Em: 488 nm/498-550 nm. Scale bar: 50 µm. Note that the detection sensitivity is different from that in Figure 2b. (**b**) Fluorescence images at 540 nm of organs dissected from mice with peritoneal dissemination of SKOV-3-luc cells, visualized with Trastuzumab-enzyme conjugates and corresponding fluorescence probes. The mice were i.p. injected with 250 µL of 1 μM Trastuzumab-Td2F2(WT), Trastuzumab-Td2F2(Mutant A) or Trastuzumab-β-galactosidase. After 3 hrs, 150 μL of 1 mM HMRef-β-D-Fuc (for Td2F2) or HMRef-β-D-Gal (for β-galactosidase) was injected. 1 hr later, the mice were sacrificed, and the mesentery was dissected out and subjected to fluorescence imaging. Ex/Em = 490 nm/515 nm-long pass. Red arrowheads show typical clumps of cancer cells. See also Supplementary Figure 8 for unmixed images and imaging results of other organs. (**c**) Quantification of fluorescence signals in Figure 4b. Cancer nodules showing clear boundaries were randomly picked up and the fluorescence signal was quantified. To evaluate the cancer/normal ratio, we also compared the ratio of fluorescence intensity between the cancer nodules and the intestines. Note that the quantification also includes tissue autofluorescence, so a value of the ratio of less than 1 does not mean the probes are hydrolyzed more in the intestines, especially for the HMRef-β-D-Fuc/Td2F2 system. \**p*<0.05, \*\**p*<0.01, Welch ANOVA test followed by the Games-Howell posthoc test. (**d**) Fluorescence images of mice with peritoneal dissemination of SKOV-3-luc cells, visualized with HMRef-β-D-Fuc and the recombinant fusion protein of anti-HER2 nanobody (2Rs15d) with Td2F2 (mutant). The mice were i.p. injected with 250 µL of 1 μM antiHER2 nanobody-Td2F2(WT/MutantA). 3 hrs later, 150 μL of 1 mM HMRef-β-D-Fuc was injected. Thirty minutes later, the mice were sacrificed and subjected to imaging after dissection. Ex/Em = 490 nm/515 nm-long pass.

Finally, taking advantage of the monomeric character of Td2F2, we also tested Td2F2 variants genetically fused to anti-HER2 nanobody 2Rs15d^37^, since the fusion protein can be purified from *E. coli* and used immediately without downstream conjugation. After confirming the function of the fusion protein for live cell imaging of SKOV-3 cells (Supplementary Figure 9), the conjugate was tested for visualization of SKOV-3 cells in the peritoneal dissemination model. As shown in Figure 4d, the cells disseminated in the peritoneal cavity were clearly visualized by HMRef-β-D-Fuc, especially with the Td2F2 Mutant A conjugate. Notably, the cancer cells in the peritoneal cavity were clearly visualized even without dissecting out the mesentery (Figure 4d), which is hard to achieve with the β-galactosidase-based system due to the insufficient bioorthogonality of the probe (Supplementary Figure 10). The successful application of the Td2F2 mutant genetically fused with an antigen-binding protein nanobody strongly suggests that it would be possible to prepare Td2F2 conjugates targeting a variety of antigens.

## Discussion

We have established a novel combination of a bioorthogonal fluorescence probe, HMRef-β-D-Fuc, and a metagenomic glycosidase, Td2F2, engineered to enhance the probe-hydrolyzing activity. By employing Td2F2 Mutant A conjugates with proteins targeting cell-surface antigens of cancer cells, vivid cancer imaging could be achieved upon administration of HMRef-β-D-Fuc with a minimal background. The exchangeability of the antigen-binding protein suggests that this system can in principle be easily adapted to visualize cells expressing any antigen of interest, though for the visualization of cancer cells, it would be necessary to confirm the specificity of the target biomarker antigen.

Cancer imaging based on antibody (or antigen-binding protein)-directed enzyme delivery is advantageous because it enables signal amplification through enzymatic turnover, in contrast to direct conjugation of the fluorophore to the antibody^14^. On the other hand, bioorthogonality is essential to minimize the background signal. Surprisingly, however, the bioorthogonality of the probes used in common reporter enzyme systems is often not sufficient, or has not been well-established. Also, the performance of a pioneering potentially bioorthogonal probe-enzyme pair has not been evaluated in detail for practical applications *in vivo*^38^. To our knowledge, the HMRef-β-D-Fuc/Td2F2 mutant system developed in this study is the first bioorthogonal probe-enzyme pair that has been shown to have sufficient activity for cancer imaging *in vivo*. Although we tested delivery of Td2F2 based on conjugation with antigen-binding proteins, delivery of Td2F2 in other ways (e.g., by viral transduction) might also be an option to achieve cancer-specific delivery. Furthermore, our highly active Td2F2-based reporter system could be employed for a variety of applications such as ELISA for biological research and diagnostics.

For the future, changing the structure of the substrate would be an interesting approach to extend the scope of our system. For example, the insertion of a self-immolative linker between the fluorophore and the sugar moiety would affect the reaction speed^39^. Further, the fluorophore could be replaced by a drug whose activity can be caged by adding the sugar moiety (i.e., forming a prodrug). Considering the bioorthogonality and high *k*_cat_/*K*_m_ values, the combination of β-D-Fuc-based prodrug and Td2F2 mutant could become a best-in-class system for various forms of DEPT, including antibody-directed enzyme prodrug therapy (ADEPT)^11^ and lectin-directed enzyme prodrug therapy (LDEPT)^40^. Another possibility would be expanding the repertoire of available enzymes. If appropriate enzymes for the other candidate bioorthogonal probes found in this study (β-L-Gal, β-D-Xyl, α-D-Ara, α-L-Ara) can be found/developed, simultaneous multi-color imaging of multiple antigens would become possible with each probe contributing a different color. Technically, we have shown that a SPiDER-type probe that becomes fluorescent and simultaneously immobilized within the cells after the enzymatic reaction enables efficient FACS-based pooled screening of the mutant activity without the need for a complicated droplet-based expression/evaluation system. If enzymes with moderate hydrolytic activity towards other bioorthogonal probes are discovered, it should be possible to enhance their activity by employing the same method. It should be noted that an important potential issue facing clinical application of the Td2F2-based system would be the immunogenicity of the enzyme, since Td2F2 is a metagenomic enzyme. However, computer-aided engineering of Td2F2 to reduce its immunogenicity could overcome this hurdle^41,42^.

Overall, we believe this study expands the chemical medicine toolbox by providing a new bioorthogonal fluorescence probe and targeted reporter enzyme pair, as well as a design strategy for engineering other candidate probe and enzyme pairs in the future.

## Supporting information

Supplementary Tables 1-2, Supplementary Scheme 1, Supplementary Figures 1-10.

## Author contributions

Z.W. and R.Kojima designed and conducted the majority of the experiments. R. Kiji contributed to the evaluation of the properties of recombinant Td2F2 and data analysis. K.F. generated the fluorescence probe library and advised on the experiments. R.T. conducted structure prediction of Td2F2 mutants and contributed to the related discussion. T.U. discovered Td2F2 as a candidate enzyme and provided the DNA template. Y.M., K.I., H.N, T.M., M.K. advised on the experiments and analysis. R. Kojima and Z.W. co-wrote the manuscript with input from all the authors. R.Kojima and Y.U. designed and supervised the entire project.

## Acknowledgment

We thank Dr. Saburo Tsuru and Dr. Chikara Furusawa for help with the FACS experiments, Mr. Naoki Seike for help with the expression of nanobody-fused Td2F2, Dr. Reiko Tsuchiya for help with the *in vivo* experiments, and Dr. Takahiro Mori for technical advice on directed evolution. This work was supported by Grants-in-Aid for Scientific Research (KAKENHI) (20H02874, 24K01642, 24H00868 to R.Kojima, 19H05632, 24H00050 to Y.U.), The Mochida Memorial Foundation for Medical and Pharmaceutical Research (grant to R.Kojima), Toyota Riken Scholarship (to R.Kojima), Nakatani Foundation Research Grant (to R.Kojima), Daiichi Sankyo Foundation for Life Science (grant to R.Kojima), Japan Science and Technology Agency (JST) PRESTO program (JPMJPR17H5 to R.Kojima), JST FOREST program (JPMJFR214N to R.Kojima), JST Mirai Program (JPMJMI24G2 to Y.U.), and JST Moonshot Research and Development Program (JPMJMS2022 to Y.U.). Z.W. was supported by a Grant-in-Aid for JSPS fellows and by the WINGS-LST program of the University of Tokyo, and R. Kiji was supported by the WINGS-LST program of the University of Tokyo.

## Data Availability

All data supporting the conclusions of this manuscript are included in the main figures and supplementary information. Additional data can be made available upon reasonable request.

### Sample Size, Statistical Analysis, and Reproducibility

We included all data in the analysis and confirmed reproducibility. No statistical method was used to predetermine sample size, which was instead chosen based on our experience and the standard practice in the field (typically 3 or more biological replicates for biochemical assays). Unless otherwise stated, all analyses in this study were performed using Excel, GraphPad Prism, or Python.

### Correspondence

Correspondence should be addressed to Ryosuke Kojima and Yasuteru Urano.

## Methods

### Materials and general information for chemical synthesis

The fluorescence probe library including HMRef-β-D-Fuc has been reported elsewhere^9^. See supporting information for the synthesis of SPiDER-β-D-Fuc. All reagents and dry solvents for synthesis were purchased from commercial suppliers (Wako Pure Chemical, Tokyo Chemical Industries, Sigma-Aldrich Japan) and were used without further purification. The composition of mixed solvents is given as volume ratio (v/v). Dimethyl sulfoxide (DMSO, fluorometric grade) for the spectrometric measurements was purchased from Dojindo. ^1^H and ^13^C nuclear magnetic resonance (NMR) spectra were recorded on an AVANCE III 400 Nanobay (Bruker, 400 MHz for ^1^H NMR, 101 MHz for ^13^C NMR). Mass spectra (MS, ESI-TOF) were measured with a MicroTOF (Bruker). High-resolution MS (HRMS) were measured using sodium formate as an external standard. Flash chromatographic purifications were performed on an automated silica gel flash column chromatography system YFLC AI-580 (Yamazen). Preparative reversed-phase HPLC using an Inertssustain C18 4.6 mm x 150 mm column (GL Science) was performed on a PU-2080 system equipped with a MD-2010 detector (JASCO) or a PU-2087 system equipped with a MD-2010 detector (JASCO). Solvent A: 99% H_2_O, 1% CH_3_CN (TFA 0.1%). B: 99% CH_3_CN and 1% H_2_O.

### Preparation of the fluorescence probes and plasmids for the expression of recombinant proteins

Construction of the plasmids is described in detail in the supplementary information. As a general subcloning method, PCR amplification was carried out by using KOD plus neo and KOD One Master Mix (TOYOBO) with a thermal cycler (Veriti 96-well Thermal Cycler, Applied Biosystems). All oligodeoxynucleotide primers were synthesized by FASMAC. Purification of PCR products from electrophoresis gel was performed by using a Monarch® DNA Gel Extraction Kit (NEB). Ligation of fragments was conducted with ligation high ver. 2 (TOYOBO). Gibson assembly was carried out by NEBuilder HiFi DNA Assembly Master Mix (NEB). Plasmids were transformed into *E. coli* DH5α competent cells (BioElegen Technology Co., Ltd.), and purified with Monarch® Miniprep Kit (NEB). Sanger sequencing was conducted by Azenta Inc.

### Cell culture

SHIN3 cells (gift from S. Imai, Nara, Japan), SKOV3 cells (American Type Culture Collection (ATCC), HTB-77), and SKOV3-luc cells (Japanese Collection of Research Bioresources (JCRB) Cell Bank, JCRB1594) were cultured in RPMI 1640 (Fujifilm Wako), containing 10% fetal bovine serum (FBS, Biosera) and 1% (v/v) penicillin and streptomycin (PS, Fujifilm Wako). NIH 3T3-HER2 cells and NIH 3T3 cells (gift from Prof. H. Kobayashi, NIH, U.S.A.) were cultured in DMEM (high glucose) (Wako), containing 10% FBS and 1% PS. Cells were cultured at 37°C in a humidified atmosphere containing 5% CO_2_.

### General information for the mouse experiments and *in vivo* fluorescence imaging of the mouse model

Female Jcl:ICR mice (6–7 weeks old) or BALB/cAJcl-nu/nu mice (6–7 weeks old) were purchased from CLEA Japan. All experimental protocols were performed in accordance with the policies of the Animal Ethics Committee of the University of Tokyo. Fluorescence images were captured with a Maestro^TM^ In-Vivo Imaging System, Ex/Em = 490 nm/ 515 nm – long pass. Exposure time was set at 500 msec. Both the stage and lighting of the equipment were set at position 1 for imaging of mice and intestines, and set at position 2 for resected organs. For quantification, fluorescence at 540 nm was extracted, and the fluorescence intensity of organs was quantified by drawing regions of interest (ROIs) with ImageJ software. Unmixing of the image was conducted with the software provided with the Maestro^TM^ In-Vivo Imaging System.

### Stock solutions of the probes

Stock solutions of all fluorescence probes were prepared at a concentration of 10 mM in DMSO and stored at-20°C until use. When required, the probes were diluted from the stock solution, resulting in assay solutions that always contained DMSO as a cosolvent (e.g., 0.01% for 1 µM and 10% for 1 mM probe).

### *In vitro* probe screening with organ lysate

The mouse organs were dissected from wild-type mice (Jcl:ICR) and washed briefly with chilled pH 7.4 PBS (Gibco), then cut into smaller pieces while keeping them on ice. The tissues were transferred to a homogenizer (lysing matrix D, MP Biomedicals) and 1 mL of T-PER tissue protein extraction reagent (Thermo Scientific) was added. Homogenizing was performed in a FastPrep-24 (MP Biomedicals). The homogenized solution was transferred into a 1.5 ml microcentrifuge tube and centrifuged at 14000 rpm for 10 min at 4°C. The supernatant was saved as whole tissue lysate. The protein concentration was determined by BCA assay and adjusted to 1 mg/mL with pH 7.4 PBS (Gibco). Then, each candidate probe was diluted to 2 μM with the same buffer. 100 μL of 1 mg/mL lysate and 100 μL of 2 μM probe were mixed in black 96-well plates. Fluorescence intensity was measured with an Envision Plate Reader for 60 minutes. Filter set: excitation: FITC (485/14 nm), emission: FITC (535/25 nm).

### Expression and purification of Td2F2 (mutants) and β-galactosidase (LacZ)

*E. coli* BL21 (DE3) harboring pZW13 (encoding 6xHis-Td2F2 in pGEX-2T, see Supplementary Information for details of the construction; the same backbone hereafter), pZW20 (6xHis-Cys-Td2F2), pZW98 (6xHis-Cys-Td2F2 Mutant A), plasmids encoding other Td2F2 mutants in the same format, or pZW19 (encoding 6xHis-β-galactosidase (LacZ) in pGEX-2T) were grown in 250 mL of Overnight Express^TM^ Instant LB Medium (Novagen), supplemented with 100 μg/mL ampicillin, at 37°C for 16∼18 hours. The cells were harvested by centrifugation at 4°C and 4000 x g for 20 min. The cell pellet was resuspended in lysis buffer with lysozyme and benzonase (QIAGEN). The crude extract was incubated on ice for 30 min. Cell debris was removed by centrifugation at 4°C and 14000 x g for 30 min. The resulting supernatant was purified using an Ni-NTA Fast Start Kit (6) (QIAGEN) to obtain the enzyme. The enzyme fraction was concentrated using an ultrafiltration membrane (Amicon Ultra, Millipore (Billerica, MA)) and the medium was replaced with 1 mL of 10 mM HEPES buffer, pH 7.4. Expression and concentration of each enzyme were checked by SDS-PAGE. For this, samples were boiled with 10% (v/v) 2-mercaptoethanol with SDS-PAGE sample buffer, and electrophoresis was performed using Mini-PROTEAN TGX Gels (Bio-Rad) at 200 V for 30 minutes. The gels were stained with Coomassie Brilliant Blue G-250 (TCI) and BCA protein assay reagent (Pierce™ BCA Protein Assay Kit (Thermo Scientific). Throughout the study, molar concentration of β-galactosidase was calculated in terms of the monomer. All samples were stored at-80°C.

### Measurement of Td2F2 and β-galactosidase activity *in vitro*

The time course of fluorescence intensity of HMRef or SPiDER was monitored with an Envision plate reader (Perkin Elmer) with a filter set of excitation = FITC (485/14 nm), emission = FITC (535/25 nm) in 10 mM HEPES buffer (pH 7.4). The fluorescence intensity changes were recorded for 30-60 minutes.

For the determination of enzyme parameters, the reaction velocity was determined at seven different substrate concentrations (ranging from approximately 0.5 to 5.0 × *K*_m_ where possible for each substrate). The *K*_m_ and *k*_cat_ were calculated by a nonlinear least-squares fitting of the kinetic data to the Michaelis-Menten equation using Kaleida Graph 4.5J (Synergy Software).

### *In vivo* evaluation of the bioorthogonality of HMRef-β-D-Fuc and HMRef-β-D-Gal

#### Evaluation of fluorescence of organs after i.p. injection of probes (Figure 1e)

BALB/c-nu/nu mice were i.p. injected with 150 μL of 1 mM fluorescence probes in PBS (containing 10 % DMSO as a cosolvent; see “Stock solutions of the probes”), and the main organs were dissected out after 30 min. Fluorescence from each organ was measured with a Maestro In-vivo Imaging System (Ex/Em = 490 nm/515 nm). Although all the images were captured under the same conditions, the brightness of the overall images varied (probably due to the unstable intensity of excitation light). Therefore, we normalized the fluorescence intensity of each image as follows: we calculated the average fluorescence intensity of each organ in each experiment set and calculated the correction ratio of the values of each experiment vs experiment 1 (for experiments 2 and 3). Then, we multiplied the raw fluorescence value and the correction ratio and averaged the results of each experiment.

#### Fluorescence imaging of peritoneal cavity after i.p injection of the probe (Figure 1f)

150 μL of 1 mM HMRef-β-D-Fuc or HMRef-β-D-Gal was i.p.-injected into Balb/c-nu/nu mice. After 1 hour, mice were sacrificed by exposure to CO_2_ gas, and the mesentery and organs were dissected out for imaging.

### Preparation of the conjugate of avidin with Td2F2 WT / β-galactosidase

Biotin-succinimidyl ester (biotin-SE, Fujifilm Wako) (20 μL of 8 mM stock solution in DMSO) was added to 2 mL of 20 μM Td2F2 in 10 mM NaPi buffer (pH 8.5, buffer exchange was done with an ultrafiltration membrane (Amicon Ultra Filter 10K 4, Millipore (Billerica, MA)) to give a final concentration of 80 μM, and the mixture was incubated for 1 hr at r.t. Then, the buffer was changed to 10 mM HEPES (pH 7.4) with an Amicon Ultra Filter to remove excess biotin-SE. After determining the concentration of the biotin-labeled enzymes with a Pierce BCA assay kit (Thermo Fisher Scientific), the avidin solution and the biotin-labeled enzyme solution were stored separately at-80°C. Equimolar amounts of avidin (as the tetramer) and biotin-enzyme (each in 10 mM HEPES buffer (pH7.4)) were mixed to generate the complex before use (see following methods for details).

### Preparation of the conjugate of Trastuzumab with Td2F2 (WT or Mutant A)/β-galactosidase (LacZ)

Trastuzumab (Herceptin; Chugai Pharmaceutical) was diluted with 10 mM HEPES (pH 7.4) to 2 mg/ml (approx. 14 μM). To prepare maleimide-labeled antibody, 3 μL of 1 mg/mL (approx. 2.3 mM) of sulfo-SMCC (Dojindo) (in 10 mM HEPES (pH7.4) containing 1 % DMSO as a cosolvent, approx. 5 eq.) was added to 100 μL antibody solution and the resulting solution was mixed and incubated at 30°C for 30 min. Excess sulfo-SMCC was removed using an Amicon ultrafilter, and the concentration of the maleimide-labeled antibody was adjusted to 1∼10 μM. To generate the antibody-enzyme conjugate, equimolar amounts of the maleimide-labeled antibody solution in 10 mM HEPES (pH 7.4) and the enzyme solution (Cys-Td2F2 [mutants] or β-galactosidase) in 10 mM HEPES (pH 7.4) were mixed. After incubation at 30°C for 40 min, the reaction was terminated by adding β-mercaptoethanol at a final concentration of 10 μM.

### Preparation of the fusion protein of anti-HER2 nanobody and Td2F2 (Mutant)

SHuffle T7 Express Competent *E. coli* (NEB) cells harboring pZW101 or pZW111 (each expressing 14xHis-bdSUMO-anti-HER2 nanobody (2Rs15d)-Td2F2 (WT or Mutant A)) were grown in 15 mL of 2x YT medium, supplemented with 100 μg/mL ampicillin, at 30°C. After the OD600 had reached 0.5-0.8, IPTG was added at a final concentration of 0.2 mM to induce expression. The cells were incubated for 20-24 hours at 18°C, then harvested by centrifugation at 4°C and 4000 x g for 20 min. The cell pellet was resuspended in Bugbuster Mastermix (Novagen). The crude extract was incubated at room temperature for 20 min. Cell debris was removed by centrifugation at 4°C and 16000 x g for 20 min. The resulting supernatant was purified using a His-spin trap (Cytiva) to obtain the fusion protein of 14xHis-bdSUMO-anti-HER2 nanobody (2Rs15d)-Td2F2 (WT or Mutant A). The enzyme fraction was concentrated and transferred to 10 mM HEPES buffer, pH 7.4, using an ultrafiltration membrane Amicon Ultra 10K (Millipore). To cleave the 14xHis-bdSUMO part, the fusion protein was treated with recombinant bdSENP1 (expressed from pSF1389 (addgene #104962)) as described by Frey et al^44^. The anti-HER2 nanobody fusion protein (2Rs15d)-Td2F2 (WT or Mutant A) was obtained in the flow-through fraction with the His-spin trap. The protein was concentrated and transferred to 10 mM HEPES buffer, pH 7.4, using an Amicon Ultra 10K ultrafiltration membrane (Millipore). Expression and concentration of each protein stock solution were checked by SDS-PAGE analysis and BCA protein assay, respectively. All samples were stored at-80°C.

### Fluorescence live-cell imaging

#### Visualization of SHIN-3 cells with avidin-Td2F2 conjugate

To generate the avidin-Td2F2 conjugate, 1 μM biotin-Td2F2 and 1 μM avidin (each in 10 mM HEPES buffer (pH 7.4)) were mixed and incubated at room temperature for 1 hour. This conjugate solution was diluted to 100 nM by adding RPMI 1640 medium (Wako) containing phenol red, 10% FBS, 1% PS.

SHIN3 cells were seeded onto Ibidi 8-well μ-slide dishes at 1.0 x 10^5^ cells/well in 250 μL of RPMI 1640 containing 10% FBS and 1% PS, and cultured for 1 day. Then, the medium was replaced with the medium containing 100 nM avidin-Td2F2 conjugate (described above), and the cells were incubated for 16 hours. The medium was washed out with PBS (1 time) and phenol red (-) RPMI 1640 medium (3 times), then replaced with 250 μL of phenol red (-) RPMI 1640 (no FBS, no PS) containing 1 μM HMRef-β-D-Fuc. The plates were incubated at 37°C. Fluorescence confocal images were obtained with TCS SP8 X (Leica) 1 hour after probe application. For the “Td2F2” condition (without avidin), 100 nM Td2F2 was used instead of avidin-Td2F2.

#### Visualization of NIH3T3-HER2 or NIH3T3 cells with Trastuzumab-Td2F2 (mutants) conjugates

NIH3T3 HER2 cells or NIH3T3 cells were seeded at the density of 1.0 x 10^5^ cells/well in 250 μL-D-MEM (Wako) containing 10% FBS and 1% PS on Ibidi eight-well μ-slide dishes and cultured for 1 day. Then, the culture medium was replaced with 250 μL of D-MEM containing 100 nM of a Trastuzumab-Td2F2 conjugate (or non-conjugated enzyme as the control), and the cells were incubated at 37°C for 16 hours. The cells were washed with PBS (1 time) and phenol red (-) D-MEM (3 times), then 250 μL of phenol red (-) D-MEM (without FBS and PS) containing 1 μM HMRef-β-D-Fuc was added. The plates were incubated at 37°C. Fluorescence confocal images were obtained with TCS SP8 X (Leica) 1 hour after probe application.

#### Visualization of SKOV-3 cells with the fusion protein of anti-HER2 nanobody and Td2F2 mutants

SKOV3 cells were seeded at 1.0 x 10^5^ cells/well in 250 μL RPMI 1640 (Wako) containing 10% FBS and 1% PS on Ibidi eight-well μ-slide dishes and cultured for 1 day. 250 μL of RPM1 1640 medium containing 100 nM anti-HER2 nanobody-Td2F2 (WT or Mutant A) was added and the cells were incubated for 3 hours. The cells were washed with phenol red (-) RPMI 1640 medium (3 times), then the medium was replaced with 250 μL of phenol red (-) RPMI 1640 (without FBS and PS) containing 1 μM HMRef-β-D-Fuc. The plates were incubated at 37°C. Fluorescence confocal images were obtained with TCS SP8 X (Leica) at 1 hour after probe application.

### *In vivo* cancer imaging of the peritoneal dissemination model in mice

#### -Construction of the model

To construct the peritoneal dissemination model, BALB/cAJcl-nu/nu mice (female, 7 week old) were intraperitoneally injected with SHIN3 cells (2 x 10^6^ cells/mouse) or SKOV3-luc cells (3 x 10^6^ cells/mouse) suspended in PBS(-). SHIN-3 model mice were used after approximately 10∼13 days, and SKOV-3-luc model mice were used after approximately 2∼3 weeks (for the SKOV-3-luc model, sufficient growth of the cancer cells was confirmed by measuring bioluminescence with an IVIS Lumina II system when necessary)

#### Imaging of SHIN-3 model with avidin-enzyme conjugate

500 μL of 500 nM avidin-Td2F2(WT) or 100 nM avidin-β-galactosidase was i.p. injected. After 16 hours, 150 μL of 1 mM HMRef-β-D-Fuc (for Td2F2) or HMRef-β-D-Gal (β-galactosidase) was i.p. injected. After 1 hour, mice were sacrificed by exposure to CO_2_ gas, and the mesentery and organs were dissected out for imaging.

#### Imaging of SKOV-3 luc model

200 μL of 250 nM Td2F2 conjugate (Trastuzumab-Td2F2 or anti-HER2 nanobody-Td2F2) was i.p. injected. After 3-16 hours (see figure legends), 150 μL of 1 mM HMRef-β-D-Fuc was i.p. injected. 30 minutes later, mice were sacrificed by exposure to CO_2_ gas and dissected for imaging. For quantification of the fluorescence intensity of cancer nodules and background tissue, regions of interest (ROIs) were drawn with ImageJ software. Twelve cancer nodules showing clear boundaries per specimen were randomly selected, and the cancer-to-normal (intestine) ratios were calculated. Welch ANOVA test followed by the Games-Howell posthoc test was performed with Python and *p* values of < 0.05 were considered statistically significant.

### Directed evolution of Td2F2

#### Construction of library of Td2F2 mutants

Template DNA encoding wild-type Td2F2 was subject to error-prone PCR by using Taq polymerase with MnCl_2_ aiming to install 4 genetic mutations in the full-length Td2F2 on average, using oZW71 (5’ ATTATCTAGAGTCGTTTTACAACGTCGTGACTGGGAA 3’), and reverse oZW73 (5’ ATTAGAATTCCTATTATTTTTGACACCAGACCAACTGGTAATGG 3’). The PCR reaction system contained 5 ng of template, 2.5 μL of 10 μM forward primer, 2.5 μL of 10 μM reverse primer, 5 μL 10x ThermoPol reaction buffer, 1 μL of 10 mM dNTPs, 5 μL of 500 μM MnCl_2_(Sigma-Aldrich), 1 μL Taq DNA polymerase (NEB), and MilliQ to make 50 μL. Initial denaturation was carried out at 95°C for 30 sec, followed by 31 cycles of 30 sec, 95°C; 30 sec, 62°C; 2 min, 68°C in a thermal cycler (Veriti 96-well Thermal Cycler, Applied Biosystems). Sanger sequencing confirmed that the median number of mutations was about 4. PCR products were kept at 4°C at the end of the 31 cycles to complete the PCR, then cleaned up by electrophoresis and collected with an NEB Gel extraction kit. The PCR amplicons coding the mutant Td2F2 were cloned into pZW94 (pET28a plasmid bearing two BsaI sites to work as the acceptor plasmid for Golden Gate assembly) by Golden Gate assembly with BsaI-HF v2 (NEB) following the protocol recommended by NEB (acceptor plasmid: 0.025 pmmol, PCR amplicon of Td2F2 mutant: 0.05 pmmol, BsaI-HF v2: 1.5 μL, T4 ligase 0.5 μL, 10x T4 ligase 2.5 μL, MilliQ up to 25 μL, incubation at 37°C for 1 hr, followed by 5 min incubation at 68°C). The reaction solution was concentrated to 6 μL with a Monarch PCR & DNA Cleanup Kit (NEB). 2 μL of the DNA solution was electroporated into NEB10beta electro competent cells (NEB), followed by culture in 500 ml LB (Kan+) medium. The plasmid library was purified with a QIAGEN Megaprep kit.

#### FACS of BL21 cells expressing highly active Td2F2 mutants

The mutant library was electroporated by a micro pulser electroporator (BIO-RAD, program Ec1) into BL21 electro competent cells (Funakoshi) and cultured in Overnight Expression LB medium (Kan+). The wild-type control (pZW95) and the negative control (pZW25, expressing mCherry) were chemically transformed into BL21 (DE3) competent cells (Nippon Gene) and cultured in Overnight Expression LB medium. 200 μL of the culture was spun down and re-suspended in 20 μL of hypotonic buffer (1/1 (v/v) PBS/ autoclaved water) and warmed at 37°C for 5 minutes. 100 μM SPiDER-β-D-Fuc solution in MilliQ was prepared and warmed at 37°C for 5 minutes. 20 μL of the prewarmed probe containing water was added to the cell suspension to give a final probe concentration of 50 μM, and the resulting cell suspension containing the probe was incubated at 37°C for 30 minutes. After spinning down the cell pellet (3000 g, 1 min), the supernatant was removed. The cell pellet was washed twice with 500 μl of PBS, and the cells were suspended in PBS. The tubes were kept on ice at 4°C for 30 min before FACS sorting.

The cells were sorted by a FACSAria III flow cytometer (Becton Dickinson) using PBS as the sheath fluid. FACS data were processed using FlowJo software 10 (TreeStar). After extracting the cells based on FSC and SSC, the cells showing bright green fluorescence were sorted (top 0.1% for the first 3 rounds and top 10% for the 4^th^ round). Sorted cells were centrifuged at 14,000 g for 30 min, and suspended in 5 μL of MilliQ. For round 1 and round 2, genes from highly fluorescent cells were amplified from the cell suspension using oZW71 and oZW73. For rounds 3 and 4, a 2-step PCR was performed to amplify the target genes (oZW88 and oZW89 for the 1^st^ PCR (cells as the templates), oZW90 and oZW91 for the 2^nd^ PCR for clean-up) using KOD plus NEO. The templates used were 5 μL of cell suspension for the 1^st^ PCR, and 1 μL of 1000-diluted 1^st^ PCR product for the 2^nd^ PCR. The amplicons were purified with a Monarch® DNA Gel Extraction Kit after each set of PCRs. The amplicons encoding the Td2F2 mutants were cloned into pZW94 by Golden Gate assembly and electroporated into NEB beta 10 electro competent cells to extract a new library for the next round of FACS screening (100 mL culture and QIAGEN Maxiprep were used for the recovery step).

#### Confirmation of the activity of individual mutants in separate culture wells

0.5 μL of the mutant library (after Round 4) was transformed to 20 μL of ECOS^TM^ SONIC Competent *E. coli* BL21(DE3) (Nippon Gene) and colonies were grown on LB plates (Kan+). Colonies were inoculated in 96-well 2.0 mL deep well plates (Corning) containing 1 mL of Overnight Express LB (Kan+) per well (1 colony/well), and the plates were incubated at 37°C for 18 hours with shaking at 1500 rpm (MBR-022R bioshaker, Titec). 100 μL of cell culture suspension was transferred to new clear, flat-bottomed 96-well plates (Corning) and cells were spun down by centrifugation at 3000 g for 5 minutes. The supernatant was discarded. The resulting cell pellet was resuspended in 20 μL of Bugbuster (Novagen) and the suspension was incubated at room temperature for 30 minutes. The cell lysate was diluted by adding 180 μL /well of PBS (pH 7.4). Cell debris was removed by centrifugation at 3000 g for 15 minutes and 2 μL aliquots of supernatant were transferred to black 96-well plates containing 98 μL /well of PBS. Finally, 100 μL of 2 μM SPiDER-β-D-Fuc was applied to each well, and the changes in fluorescence was recorded for 30 minutes with an Envision plate reader (PerkinElmer). Plasmids encoding mutants with high activity were subjected to Sanger sequencing (Azenta).

### QM/MM calculation

For the initial protein structure, the crystal structure (PDBID: 3WH7) was used. The crystal structure was processed with PDB2PQR server 58) to assign H atoms at pH 7.4. The structure of the substrate was introduced into the protein so that its-D-fucose moiety coordinates matched the native structure in the crystal. The mutated amino acid residues were changed manually. The protein-substrate hybrid structures were optimized by a 2-layer ONIOM method with Gaussian16^45^). The calculations were performed at the level of ONIOM(B3LYP/6-31G(d):UFF) using electronic embedding. The high layer included the substrate and the side chains of amino acids (166, 296, 325, 352, 406). For the low layer, atomic charges of the substrate and the catalyst were calculated at the same level as the high layer. In the structural optimization, only atoms within 8 Å of the high layer atoms were allowed to relax.

