## Supplementary Tables 1-2, Supplementary Scheme 1, Supplementary Figures 1-10. for "Low-Background Cancer Imaging With a Bioorthogonal Fluorescence Probe and Engineered Reporter Enzyme Bearing a Targeting Moiety"

**Inventory**

**Supplementary Table 1.** Plasmids used in this study, and sequences of Td2F2 mutants ……………………….…..3
**Supplementary Table 2.** Oligonucleotides used in this study……………………………………………….…......5
**Supplementary Scheme 1.** Synthesis of SPiDER-β-D-Fuc……………………………………………..…........6
**Supplementary Figure 1.** Confirmation of expression and purification of recombinant Td2F2……………..….…..8
**Supplementary Figure 2.** Images of organs obtained in the in vivo evaluation of HMRef-β-D-Fuc and HMRef-β-D-Gal (related to Figure 1d).……………………………………………………………………………….….9
**Supplementary Figure 3.** Confirmation of the low reactivity of HMRef-β-D-Fuc with human serum.………..…....9
**Supplementary Figure 4.** Confirmation of labelability of β-galactosidase and Cys-Td2F2 with maleimide..……..10
**Supplementary Figure 5.** Analysis of a mixture of BL21(DE3) cells expressing Td2F2 WT and BL21(DE3) cells expressing mCherry, treated with either HMRef-β-D-Fuc or SPiDER-β-D-Fuc……………………………….…11
**Supplementary Figure 6.** Assay results of mutants shown in Figure 3c with SPiDER-β-D-Fuc..………………......12
**Supplementary Figure 7.** Structural analysis of the complex of Td2F2 and HMRef-β-D-Fuc by a QM/MM method…………………………………………………………………………………………...................13
**Supplementary Figure 8.** Fluorescence images of all the dissected organs in the experiment shown in Figure 4c, together with the unmixed images.……………………………………………………………………………..14
**Supplementary Figure 9.** Validation of the fusion protein of anti-HER2 nanobody (2Rs15d) and Td2F2 (Mutant A) in vitro and in cellulo.……..…………………………………………..…………………………….......15
**Supplementary Figure 10.** Fluorescence imaging of a mouse with intraperitoneal dissemination of SKOV-3 cells using the β-galactosidase system.…………………………………………………………………………....16

**Supplementary references**……………………………………………………………..…………………… .17

| Plasmids | Description and Cloning Strategy | Source |
|---|---|---|
| **pJTtd2f2** | Plasmid encoding Td2F2 WT | Uchiyama et al[1] |
| **pCMV-LacZ** | Constitutive β-galactosidase (LacZ) expression vector | Clontech |
| **pGEX-2T** | A bacterial vector to induce Tac-promoter-driven expression of GST fusion proteins with a thrombin site. | GE HealthCare |
| **pET28a** | A backbone plasmid to induce T7-promoter-driven expression pf N-terminally 6xHis-tagged proteins with a thrombin site. Addgene 51139 encodes SpySrt A, so we replaced the encoding gene. | Addgene 51139[2] |
| **pZW13** | A bacterial expression vector of 6xHis-(GGGS)-Td2F2. Insert 1 encoding 6xHis-GGGS was generated by annealing oZWan01 and oZWan02, and insert 2 encoding WT Td2F2 was PCR-amplified from pJTtd2f2 with oZW52 and oZW05b. pGEX-2T was digested with *EcoN*I/*EcoR*I-HF as a backbone, and combined with the two inserts by Gibson assembly. | This work |
| **pZW19** | A bacterial expression vector of 6xHis-(GGGS)-β-galactosidase (LacZ). LacZ was PCR-amplified from pCMV-LacZ using oZW31 and oZW32, digested with *Xba*I/*EcoR*I-HF and cloned into the corresponding sites (*Xba*I/*EcoR*I ) of pZW13. | This work |
| **pZW20** | A bacterial expression vector of 6xHis-(GGGS)-CysTd2F2. Td2F2 bearing Cys at the N-terminus was PCR-amplified from pJTtd2f2 using oZW36 and oZW07, digested with *Xba*I/*EcoR*I-HF and cloned into the corresponding sites of pZW13. | This work |
| **pZW94** | Acceptor vector for BsaI-based Golden Gate assembly (GGA), which expresses Td2F2 mutants in *E. coli* after GGA of PCR amplicons encoding Td2F2 mutants. The acceptor sequence was PCR-amplified from pU6-pegRNA-GG-acceptor (Addgene 132777[3], encoding acceptor sequences for BsaI-based Golden Gate assembly; the mRFP1 sequence is flanked by two BsaI-sites, so the success of GGA can be monitored by colony color) using oZW69 and oZW70 cloned into pET28a digested with *BamH*I/*Xho*I with Gibson assembly. | This work |
| **pZW95** | A bacterial expression vector of 6xHis-Td2F2 WT used as the wild-type control in FACS screening. Td2F2 WT was PCR-amplified from pZW13 using oZW71 and oZW73 and cloned into pZW94 by Golden Gate assembly with the aid of digestion by *Bsa*I. | This work |
| **pZW98** | A bacterial expression vector of 6xHis-Cys-Td2F2 mutant A. Cys-Td2F2 mutant A was PCR-amplified from cell lysates used for the assay in individual wells (after FACS screening) using oZW79 and oZW73, and cloned into pZW94 by Golden Gate assembly. | This work |
| **pZW25** | A bacterial expression vector of 6x-His-mCherry. mCherry was digested from pAAV-CaMKIIa-mCherry (addgene 114469) with *BamH*I/*EcoR*I-HF and cloned into the corresponding sites (*BamH*I/EcoRI ) of pZW20. | Addgene 114469 |
| **pZW101** | Bacterial expression vector of 14xHis-bdSUMO-anti HER2 nanobody 2Rs15d-Td2F2 WT. This plasmid was constructed by stepwise normal subcloning using Gibson assembly. The backbone was pHS0806 His14-bdSUMO-AwaIscB (addgene 176539[4]), and the 2Rs15d coding sequence was | This work |

| | | |
|---|---|---|
| | amplified from gene fragment synthesized by Twist Bioscience. The insert Td2F2 WT was PCR-amplified from pZW95R-WT. The sequence of the protein was as follows. (bdSUMO, 2Rs15d)<br>MSKHHHHSGHHHTGHHHHSGSHHHTGSAAGGEEDKKPAGGEGGGAHINLKVKGQDGNE<br>VFFRIKRSTQLKKLMNAYCDRQSVDMTAIAFLFDGRRLRAEQTPDELEMEDGDEIDAMLH<br>QTGGSSGSQVQLQESGGGSVQAGGSLKLTCAASGYIFNSCGMGWYRQSPGRERELVSRISG<br>DGDTWHKESVKGRFTISQDNVKKTLYLQMNSLKPEDTAVYFCAVCYNLETYWGQGTQVT<br>VSSGTLEGGGGSGGGGS-(then Td2F2, starting from A (deleting the first M) | |
| **pZW111** | Bacterial expression vector of 14xHis-bdSUMO-anti HER2 nanobody 2Rs15d-Td2F2 mutant A. The method of construction is basically the same as that for pZW101 (changing the template) | This work |

**Amino acid sequence of Td2F2 wild type**

MAGERFPADFVWGAATAAYQIEGAVREDGRGVSIWDTFSHTPGKIADGTTGDVACDSYHRYGEDIGLLNALGMNAYRFSIAWPRIVPLGA
GPINQAGLDHYSRMVDALLGAGLQPFVTLYHWDLPQPLEDRLGWGSRATATVFAEYADIVVRQLGDRVTHWATLNEPWCSAMLGYYLGVH
APGHTDLKRGLEASHNLLLGHGLAVQAMRAAAPQPLQIGIVLNLTPTYPASDSPEDVAAARRFDGFVNRWFLDPLAGRGYPQDMLDYYGA
AAPQANPEDLTQIAAPLDWLGVNYYERMRAVDAPDASLPQAQRLDDPDLPHTADREVYPEGLYDILLRLHNDYPFRPLYITENGCALHDE
IAEDGGIHDGQRQAFFEAHLAQLQRALAAGVPLKGYFAWSLLDNFEWAMGLSMRYGICYTNFETLERRIKDSGYWLRDFIAGQRG*

**Amino acid sequence of Mutant A (E27G, I24T, F243L, E296G. Mutation sites were noted by red.)**

MAGERFPADFVWGAATAAYQIEGAVRGDGRGVSTWDTFSHTPGKIADGTTGDVACDSYHRYGEDIGLLNALGMNAYRFSIAWPRIVPLGA
GPINQAGLDHYSRMVDALLGAGLQPFVTLYHWDLPQPLEDRLGWGSRATATVFAEYADIVVRQLGDRVTHWATLNEPWCSAMLGYYLGVH
APGHTDLKRGLEASHNLLLGHGLAVQAMRAAAPQPLQIGIVLNLTPTYPASDSPEDVAAARRLDGFVNRWFLDPLAGRGYPQDMLDYYGA
AAPQANPEDLTQIAAPLDWLGVNYYGRMRAVDAPDASLPQAQRLDDPDLPHTADREVYPEGLYDILLRLHNDYPFRPLYITENGCALHDE
IAEDGGIHDGQRQAFFEAHLAQLQRALAAGVPLKGYFAWSLLDNFEWAMGLSMRYGICYTNFETLERRIKDSGYWLRDFIAGQRG*

**Supplementary Table S1**: Plasmids used in this study, and sequences of Td2F2 mutants.

| Primers | Sequence |
|---|---|
| **oZWan01** | CAGTATTCATGTCCCCTATACTAGGTCACCACCACCACCACCACGGATCCGGAGAAAATCTGTATTTTCAGGGCGGTGGCGGTTCTGGCGGTGGCGGTTCTGGCGGTGGCGGTTCTGGCGGTGGCGGTTCTAGAATGGCAGGAGAACGATTCCCGGCT |
| **oZWan02** | AGCCGGGAATCGTTCTCCTGCCATTCTAGAACCGCCACCGCCAGAACCGCCACCGCCAGAACCGCCACCGCCAGAACCGCCACCGCCCTGAAAATACAGATTTTCTCCGGATCCGTGGTGGTGGTGGTGGTGACCTAGTATAGGGGACATGAATACTG |
| **oZW05b** | TCAGTCAGTCACGATGAATTCCTAGCCGCGCTGTCCGGCGATGAAGTC |
| **oZW07** | ATTAGAATTCCTAGCCGCGCTGTCCGGCGATGAA |
| **oZW31** | ATTATCTAGAGTCGTTTTACAACGTCGTGACTGGGAA |
| **oZW32** | ATTAGAATTCCTATTATTTTTGACACCAGACCAACTGGTAATGG |
| **oZW36** | ATTATCTAGATGCATGGCAGGAGAACGATTCCCGGCT |
| **oZW52** | ATGGCAGGAGAACGATTCCCGGCTGAC |
| **oZW69** | ATTAGGATCCGAGACCGTAGTCAAAAGCCTCCGG |
| **oZW70** | ATTACTCGAGGGAGACCTCCCTATCAGTGATAGAGATTGACATCCCTATCAGTGATAGAGATACTGAGCACGGATCTGA |
| **oZW71** | GGCTACGGTCTCCGATCCATGGCAGGAGAACGATTCCCGGCTG |
| **oZW73** | GGCTACGGTCTCTCGAGTCAGCCGCGCTGTCCGGCGATGAAG |
| **oZW79** | GTGCTCCACAAGAGGCGCTATGATCTGCAAGCGGTACACTCAGATCGCAAGGATTAGTTATTCATTATCAGCCGCGCTGT |
| **oZW88** | CCCGGCTGACTTTGTCTGGGGC |
| **oZW89** | AAGTCGCGCAGCCAGTAGCCAC |
| **oZW90** | GGCTACGGTCTCCGATCCATGGCAGGAGAACGATTCCCGGCTGACTTTGTCTGGGGC |
| **oZW91** | GGCTACGGTCTCTCGAGTCAGCCGCGCTGTCCGGCGATGAAGTCGCGCAGCCAGTAGCCAC |

**Supplementary Table S2:** Oligonucleotides used in this study.

### Synthesis of SPiDER-β-D-Fuc

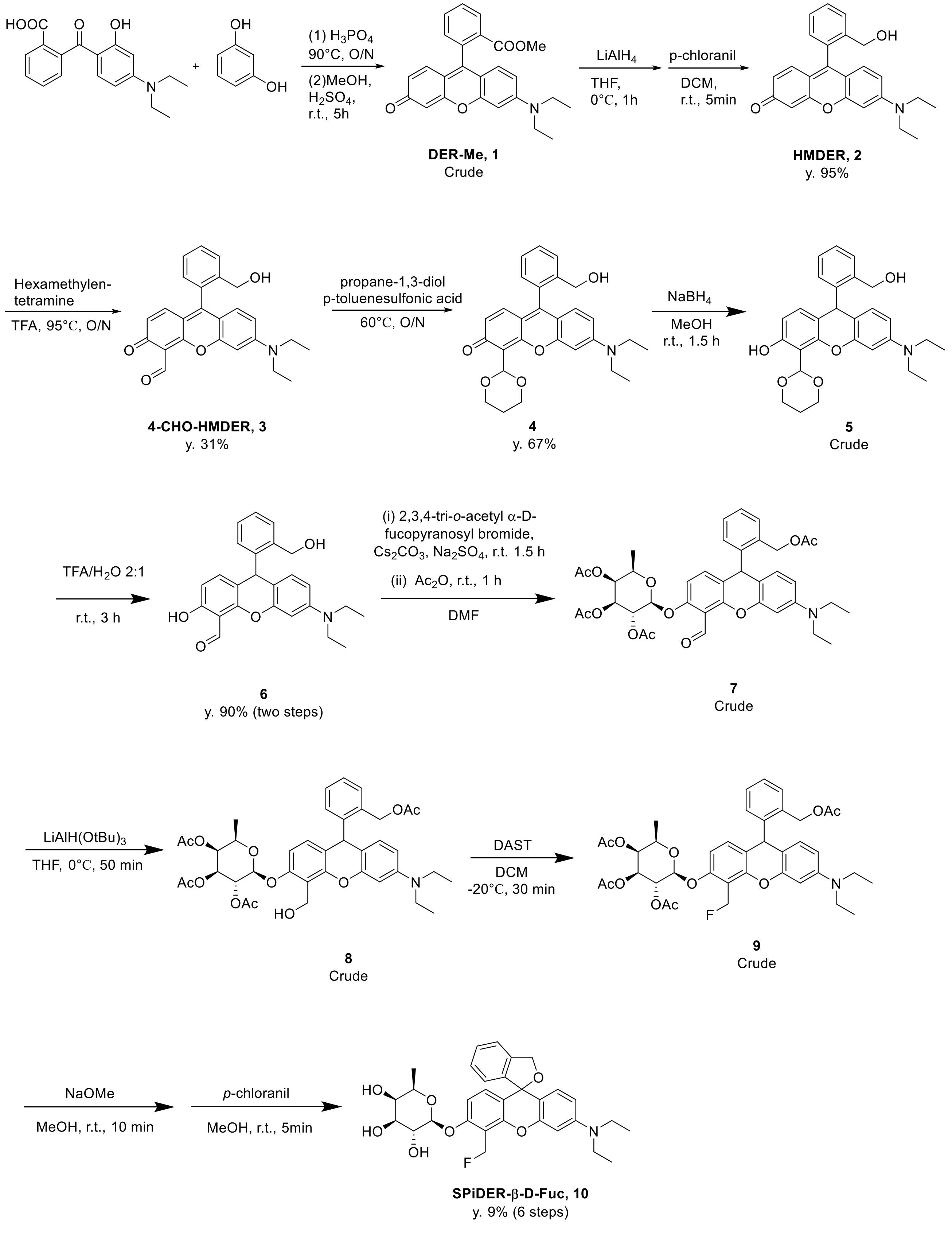


**Supplementary Scheme 1.** Synthesis of SPiDER-β-D-Fuc.

**Synthesis of Compounds 1-6**

Synthesized as described by Doura et al[5].

**Synthesis of compounds 7-10**

2,3,4-Tri-O-acetyl-α-D-fucopyranosyl bromide (3.94 g, 11.2 mmol), $Cs_2CO_3$ (5.9 g, 16.7 mmol), $Na_2SO_4$ (1.5 g, 10.6 mmol) and compound **6** (900 mg, 2.2 mmol) were added to dry DMF (20 mL) and the mixture was stirred at room temperature for 2.5 hours. Then acetic anhydride (324 mg, 3.0 mmol) was added and stirring was continued at room temperature for 1 hour. After evaporation of DMF, saturated $NH_4Cl$ aq. was added to neutralize the solution. The resulting mixture was extracted with $CH_2Cl_2$ (3 times), and the combined organic phase was dried over $Na_2SO_4$, filtered through Celite, and evaporated. The residue was purified by flash chromatography ($CH_2Cl_2$/EtOAc = 98/2) to obtain crude compound **7** as a pale pink solid. HRMS-ESI (m/z): $[M + Na]^+$ calculated for 740.2686 ($C_{39}H_{43}NO_{12}Na$), found 740.2678.

Next, 1.0 M $LiAlH(OtBu)_3$ in THF (2 mL) and crude **7** (950.9 mg) were dissolved in dry THF (8 mL) at 0 °C, and the mixture was stirred at 0 °C for 50 minutes. Saturated $NH_4Cl$ a.q. (6 mL) was added to the solution on ice to quench the reaction. EtOAc (14 mL) was added and the whole was stirred at room temperature for 1 hour, then the organic phase was separated. $CH_2Cl_2$ and saturated potassium sodium tartrate aqueous solution were added to the aqueous phase. The reaction mixture was stirred, and the organic phase was separated (2 times). The combined organic phase was evaporated, and the residue was roughly purified by flash chromatography ($CH_2Cl_2$/EtOAc = 98/2) to obtain crude **8** as a pale pink solid. HRMS-ESI (m/z): $[M + Na]^+$ calculated for 742.2835 ($C_{39}H_{45}NO_{12}Na$), found 742.2834.

DAST (400 μL, 488 mg, 3.02 mmol) and crude **8** were dissolved in dry $CH_2Cl_2$ (10 mL) and the solution was stirred at room temperature for 30 minutes. MeOH (20 mL) was added on ice to quench the reaction and the mixture was evaporated. The residue (containing compound **9**) and NaOMe (1 g, 18.51 mmol) were dissolved in MeOH (20 mL) and the solution was stirred at room temperature for 5 minutes. After evaporation of the solvent, saturated $NH_4Cl$ a.q. (5 mL) was added to neutralize the residue. Saturated $NaHCO_3$ aq. was added, and the mixture was extracted with $CH_2Cl_2$ (3 times). The combined organic phase was dried over $Na_2SO_4$, filtered through Celite, and evaporated. The residue was dissolved in MeOH (20 mL), then NaOMe (200 mg, 3.70 mmol) was added, and the mixture was stirred at room temperature for 10 minutes. p-Chloranil (200 mg, 0.81 mmol) was added and stirring was continued at room temperature for 5 minutes. Saturated $NH_4Cl$ aq. (1 mL) was added to quench the reaction, and the resulting solution was evaporated. Saturated $NaHCO_3$ aq. was added to the residue, and the mixture was extracted with $CH_2Cl_2$ (3 times). The combined organic phase was washed with brine, filtered through Celite, and evaporated. The residue was purified by flash chromatography ($CH_2Cl_2$/MeOH = 98/2 to 94/6) followed by preparative reverse-phase HPLC (A/B = 50/50, 2 times). The eluate was evaporated and extracted with $CH_2Cl_2$ (3 times). The combined organic phase was evaporated and dried to obtain pure compound **10** (SPiDER-β-D-Fuc) as a pale pink solid (114.6 mg, y. 9% in 6 steps).

$^1H$ NMR ($CD_3CN$): δ. 7.46 (d, 1H, J = 7.5 Hz), 7.40 (t, 1H, J = 7.3 Hz), 7.29 (t, 1H, J = 7.3 Hz), 6.98 (dd, 1H, J = 2.6 and 8.9 Hz), 6.90 (t, 1H, J = 4.2Hz), 6.86 (q, 1H, J = 5.8 Hz), 6.76 (d, 1H, J = 8.8Hz), 6.54 (d, 1H, J = 2.5Hz), 6.49 (dd, 1H, J = 2.6 and 8.8 Hz), 5.83 (d, 2H, J = 48.2 Hz), 5.28 (s,2H), 4.88 (dd, 1H, J = 7.6 and 18.0 Hz), 3.74-3.53 (m, 4H), 3.40 (q, 4H, J =7.0 Hz), 2.19 (s, 3H), 1.16 (t, 6H, J = 7.0 Hz).

$^{13}C$ NMR ($CD_3CN$): δ. 156.3, 150.9, 148.5, 145.1,139.0, 138.9, 131.0, 129.1, 127.9, 127.6, 122.9, 120.7, 111.0,109.8, 108.2, 100.8, 100.4, 96.8, 82.9, 74.0, 73.1, 72.3, 70.7, 70.3, 43.7, 40.0, 15.3, 11.5.

HRMS-ESI (m/z): $[M + H]^+$ calculated for 552.2392 ($C_{31}H_{35}FNO_7$), found 552.2387.

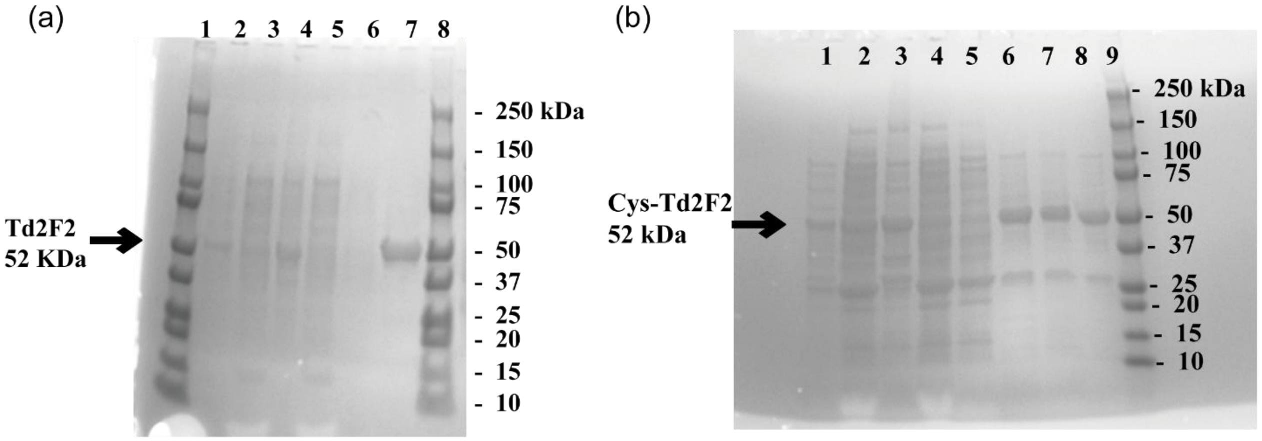


**Supplementary Figure 1**. Confirmation of expression and purification of Td2F2. (**a**) SDS-PAGE analysis of His-tagged Td2F2 expressed by BL21 (DE3) cells using the Overnight Express™ system and purified on an Ni-NTA column. Lane 1: molecular markers, lane 2: whole cell lysate, lane 3: supernatant after centrifugation of the cell lysate, lane 4: pellet, lane 5: flow-through fraction, lane 6: wash fraction, lane 7: eluted fraction; lane 8, molecular markers. (**b**) SDS-PAGE analysis of His-tagged Cys-Td2F2 expressed by BL21 (DE3) cells using the Overnight Express™ system and purified on an Ni-NTA column. Lane 1, whole solution; lane 2, supernatant; lane 3, pellet; lane 4, flow-through fraction; lane 5, wash fraction; lane 6, eluted fraction 1; lane 7, eluted fraction 2; lane 8, solution after desalting; lane 9, molecular markers. For both (**a**) and (**b**), the gels were stained with CBB before observation.

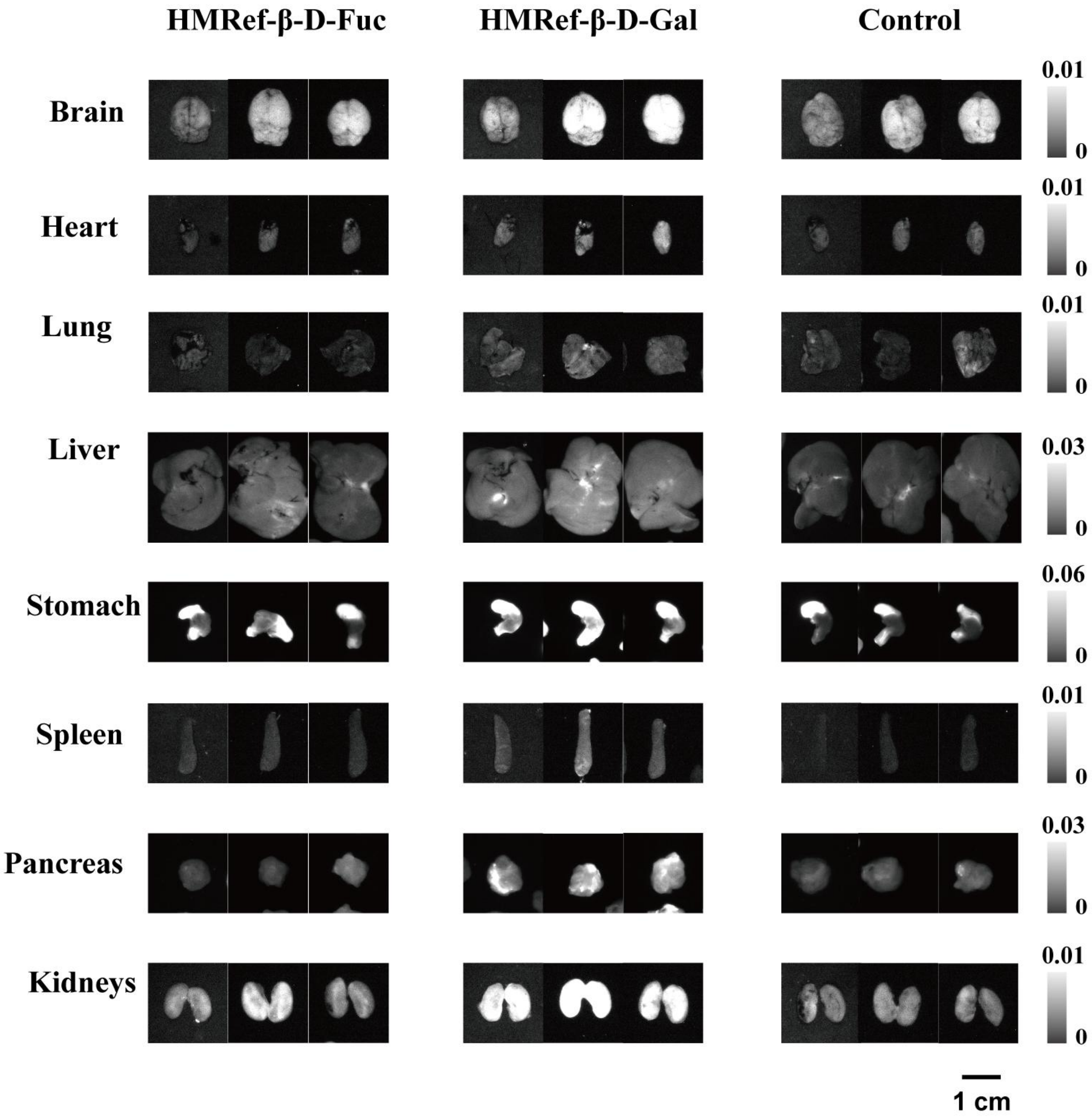


**Supplementary Figure 2**. Images of organs obtained in the in vivo evaluation of HMRef-β-D-Fuc and HMRef-β-D-Gal (related to Figure 1e). Fluorescence images at 540 nm of organs from BALB/c-nu/nu mice i.p. injected with 150 μL of 1 mM fluorescence probes and dissected at 30 minutes after injection. Fluorescence from each organ was measured with a Maestro In-vivo Imaging System (Ex/Em = 490 nm/515 nm). Note that the fluorescence includes tissue autofluorescence (we did not conduct unmixing because it would have made quantification impossible).

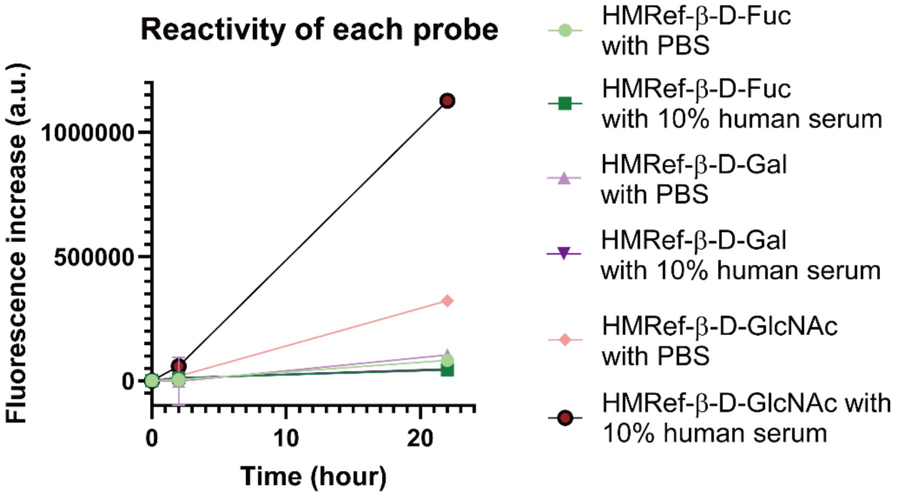


**Supplementary Figure 3**. Confirmation of the low reactivity of HMRef-β-D-Fuc with human serum. 1 μM of each probe was incubated either in PBS(-) or PBS (-) containing 10 % human serum (Jackson ImmunoResearch Lab. Inc), and the fluorescence increase was monitored with an Envision plate reader.

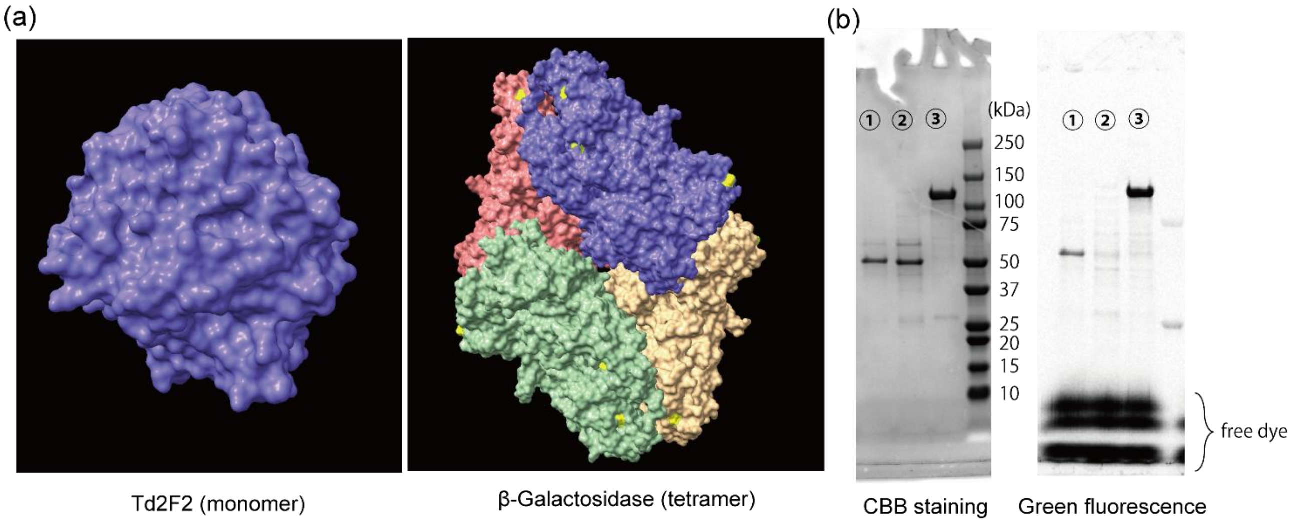


**Supplementary Figure 4**. Confirmation of labelability of β-galactosidase and Cys-Td2F2 with maleimide. (**a**) 3D structure of β-galactosidase (LacZ, tetramer) (PDB ID: 1JYN) and Td2F2 (PDB ID: 3WH5). Thiols of Cys residues are highlighted in yellow. One subunit of β-galactosidase possesses 12 cysteines, and several are exposed at the surface. Each Td2F2 monomer contains 4 cysteines, but none of them are exposed. Therefore, cysteine was artificially added at the N-terminus of Td2F2 for bioconjugation. (**b**) Confirmation of the labelling of the recombinant proteins expressed from BL21 (DE3) cells. 2.5 μM of each enzyme (① Cys-Td2F2 ② Td2F2 ③ β-galactosidase (LacZ)) in 100 mM NaPi buffer (pH 6.4, containing 150 mM NaCl, 25 μM TCEP, and 10% (v/v) of glycerol) was reacted with 10 eq of fluorescein-5-maleimide at 37ºC for 2 hours. The reaction was quenched by adding an excess of β-mercaptoethanol, and the solution was subjected to SDS-PAGE. Left: CBB staining to show total protein. Right: Fluorescence image of the gel (iBright FL1500, filter set for Alexa Fluor 488). The results show that Cys installation is necessary for labelling Td2F2, but not for labelling β-galactosidase. Thus, site-specific labelling can be achieved in the case of Cys-Td2F2.

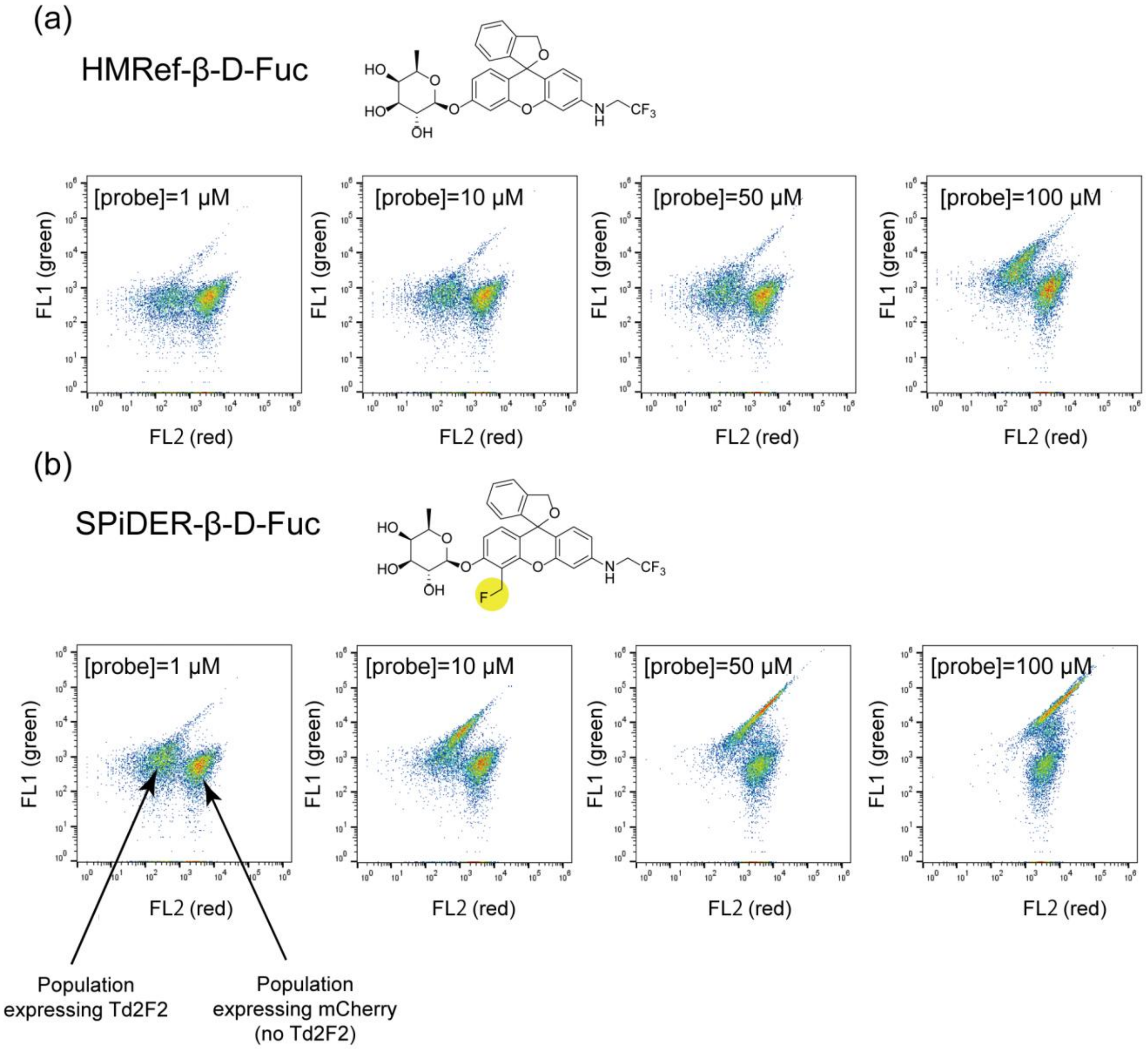


**Supplementary Figure 5**. Analysis of the mixture of BL21(DE3) cells expressing Td2F2 WT and BL21(DE3) cells expressing mCherry, treated with either HMRef-β-D-Fuc (**a**) or SPiDER-β-D-Fuc (**b**) for 30 min. Y axis (FL1): green, X axis (FL2): red. When cells were treated with HMRef-β-D-Fuc, green fluorescence of both Td2F2$^+$ and mCherry$^+$ increased, indicating that the fluorescent product (HMRef) had leaked out from the Td2F2$^+$ cells, thus failing to label the Td2F2-expressing cells with single-cell resolution. When cells were treated with SPiDER-β-D-Fuc, green fluorescence of Td2F2$^+$ cells drastically increased while that of mCherry$^+$ did not, even in the mixed culture, indicating that the Td2F2 activity can be detected at a single-cell resolution in this case. Note that the red fluorescence of Td2F2$^+$ cells also increased upon the probe treatment, possibly due to the change of cellular conditions caused by the reaction with quinone methide. We chose FACS screening because the activity of Td2F2 was evidently captured with single-cell resolution.

(a)

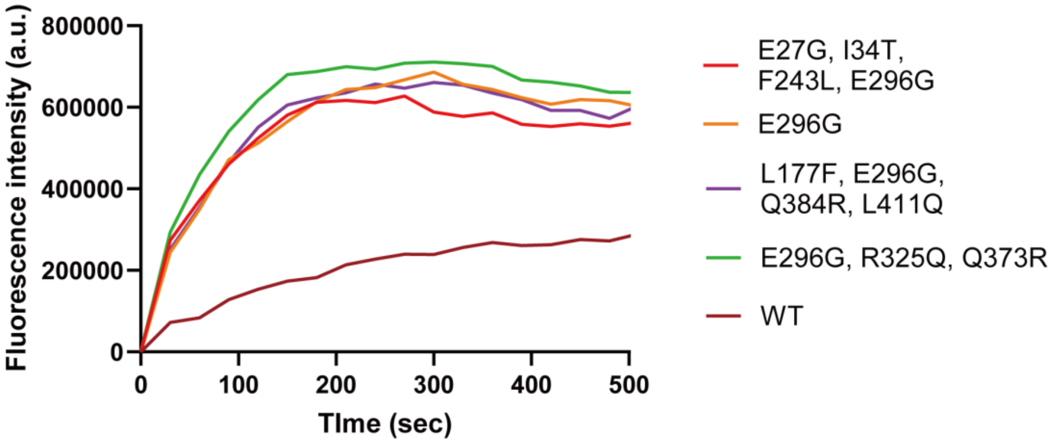


(b)

| | $K_m$ (μM) | $k_{cat}$ (/s) | $k_{cat}/K_m$ (/M/s) | ratio |
|---|---|---|---|---|
| Td2F2 WT | 21.4 | 4.2 | $2.0 \times 10^5$ | 1.0 |
| Td2F2 mutant A | 2.8 | 4.6 | $1.6 \times 10^6$ | 8.4 |

**Supplementary Figure 6.** Assay results of mutants shown in Figure 3c with SPiDER-β-D-Fuc. (**a**) The assay results of individual mutants in the homogeneous solution, using the same procedure as for Figure 3c. (**b**) Enzymatic parameters of Td2F2 WT and Mutant A (E29G, I34T, F243L, E296G), using SPiDER-β-D-Fuc as a substrate.

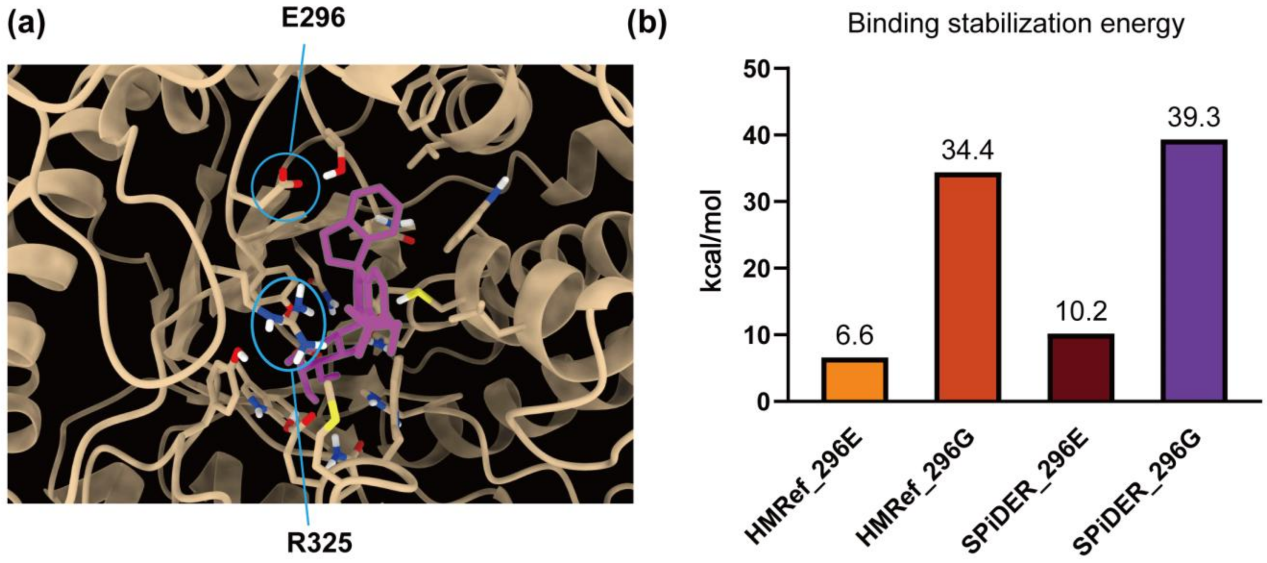


**Supplementary Figure 7**. Structural analysis of the complex of Td2F2 and HMRef-β-D-Fuc by a QM/MM method. (**a**) Calculated structure of Td2F2 (WT) with HMRef-β-D-Fuc. The 296$^{th}$ glutamate (E) and the 325$^{th}$ Arg (R) are highlighted. It seems that the E296G mutation structurally facilitates the probe's access to the enzyme's active site. Additionally, E296 forms a salt bridge with R325, and the E296G mutation may eliminate this interaction, further improving accessibility. (**b**) Calculated binding stabilization energies of Td2F2 (WT or E296G) and HMRef-β-D-Fuc/ SPiDER-β-D-Fuc. The calculation suggests that the E296G mutation significantly contributes to stabilization of the encounter complex of the enzyme and the substrate, in accordance with the experimental finding that the E296G mutant shows much lower $K_m$ values for substrates, compared to WT. See Supplementary Data 1-6 for the other calculated structures.

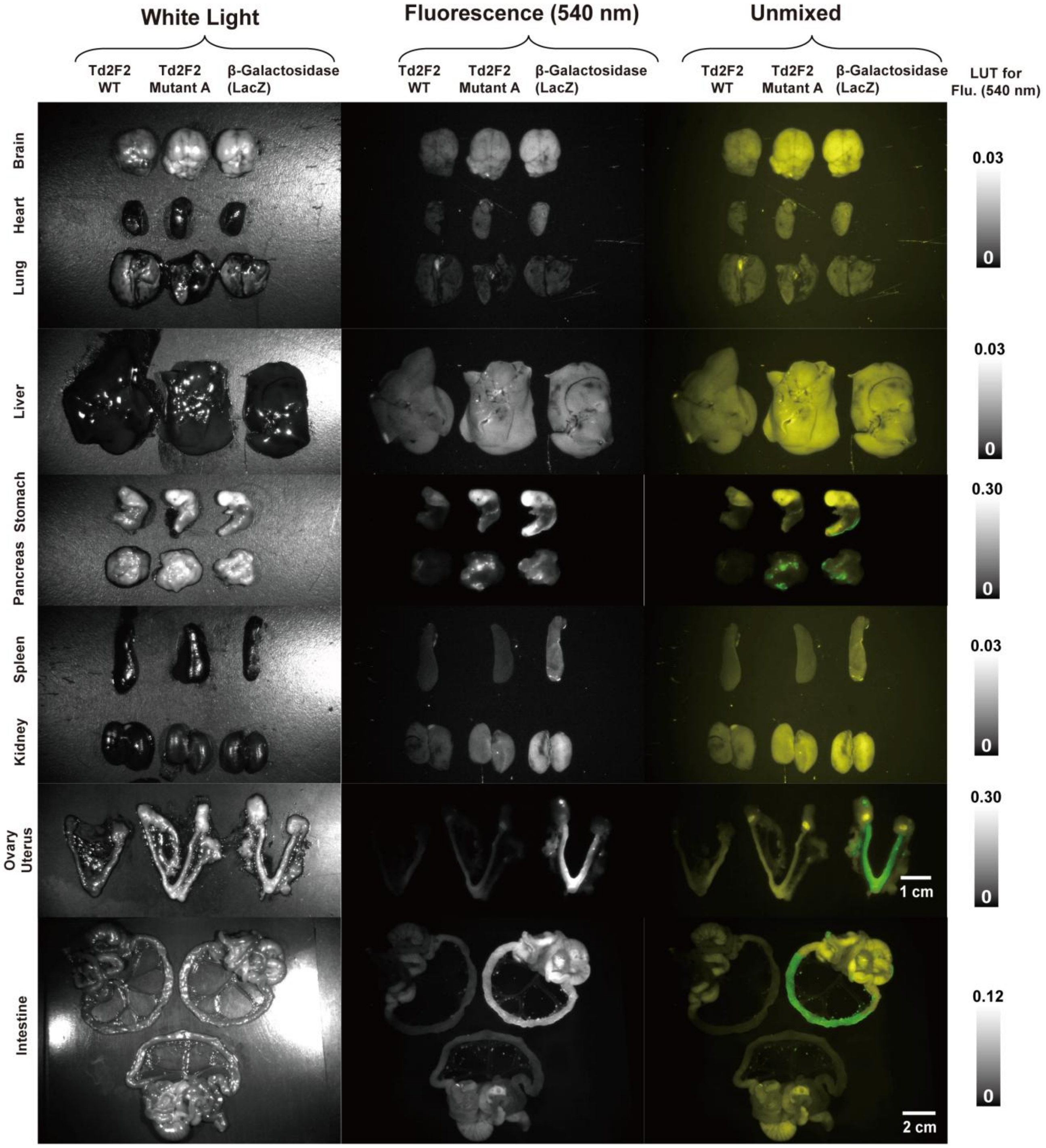


**Supplementary Figure 8**. Fluorescence images of all the dissected organs in the experiment shown in Figure 4c, together with the unmixed images. The images of "Intestine" are the same as those in Figure 4c. LUT: look up table. BALB/c-nu/nu mice were i.p. injected with 150 μL of 1 mM fluorescence probes, and the main organs were dissected out after 30 min. Fluorescence from each organ was measured with a Maestro In-vivo Imaging System (Ex/Em = 490 nm/515 nm). Fluorescence at 540 nm was extracted. Unmixed images were constructed by the software supplied with the Maestro In Vivo imaging system.

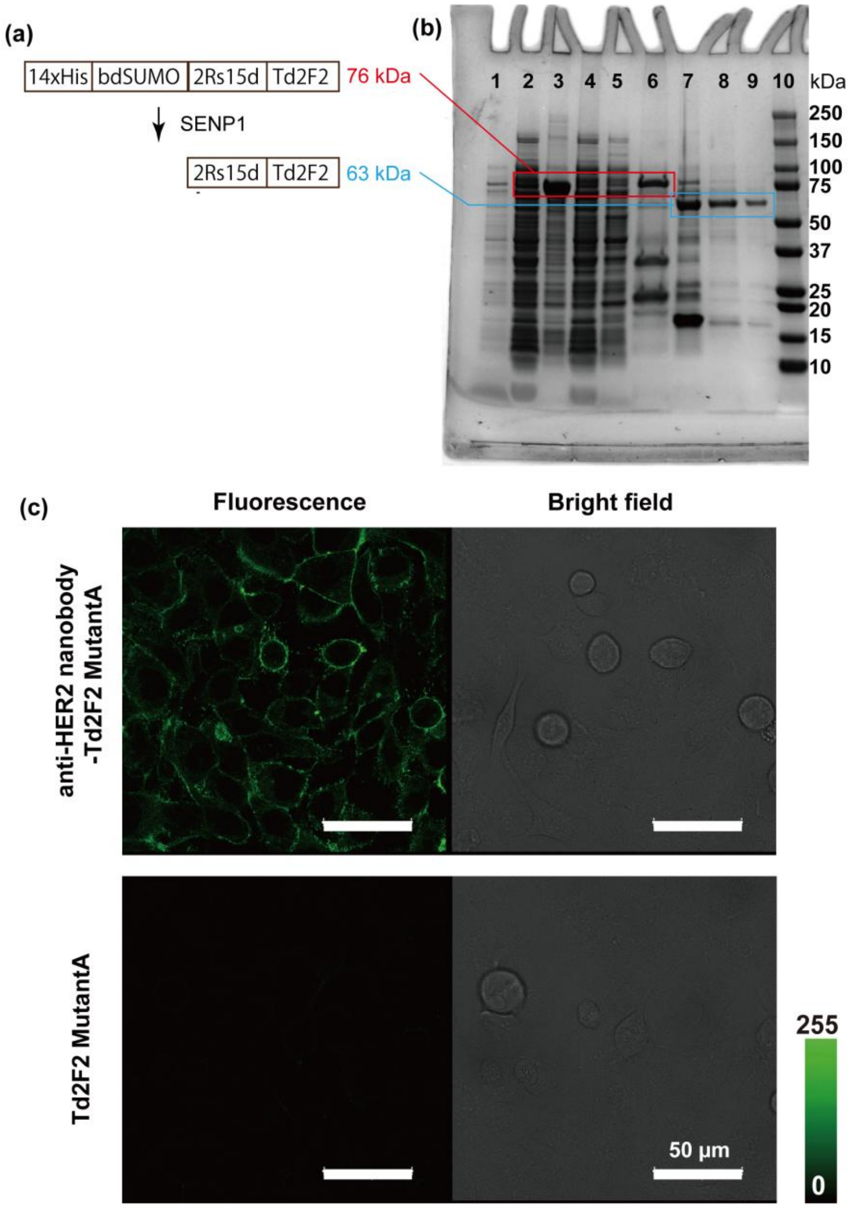


**Supplementary Figure 9.** Validation of the fusion protein of anti-HER2 nanobody (2Rs15d) and Td2F2 (Mutant A) in vitro and in cellulo. (**a**) Schematic illustration of the expression and purification of anti-HER2nanobody-Td2F2. 14xHis-bdSUMO-2Rs15d-Td2F2(Mutant A) was purified on Ni-NTA resin via the His tag. Then, 14x-His-bdSUMO was cleaved by SENP1, and the fusion protein 2Rs15d-Td2F2 was obtained in the flow-through fraction of the 2nd His-tag purification. (**b**) SDS-PAGE image to confirm the purity of 2Rs15d-Td2F2(Mutant A). **1**, whole solution after protein expression in Shuffle T7 Competent *E. coli*; **2**, supernatant after lysis; **3**, pellet after lysis; **4**, flow-through fraction of 1st His-tag purification; **5**, wash fraction of 1st His-tag purification; **6**, eluted fraction of 1st His-tag purification (14xHis-bdSUMO-2Rs15d-Td2F2(Mutant A)); **7**, product of SENP1 cleavage reaction; **8**, flow-through fraction of 2nd His-tag purification after SENP1 reaction (containing 2Rs15d-Td2F2); **9**, final product; **10** molecular markers. (**c**) Fluorescence confocal images of SKOV3 cells treated with 100 nM anti-HER2nanobody(2Rs15d)-Td2F2(Mutant A) or Td2F2(Mutant A) without the antigen-recognition moiety. The cells were treated with enzyme in the medium for 16 hours. After washing 3 times, 1 μM HMRef-β-D-Fuc was applied for 1 hour. Fluorescence images were obtained with TCS-SP8. HyD, Ex/Em = 488 nm / 498-550 nm. Scale bars: 50 μm.

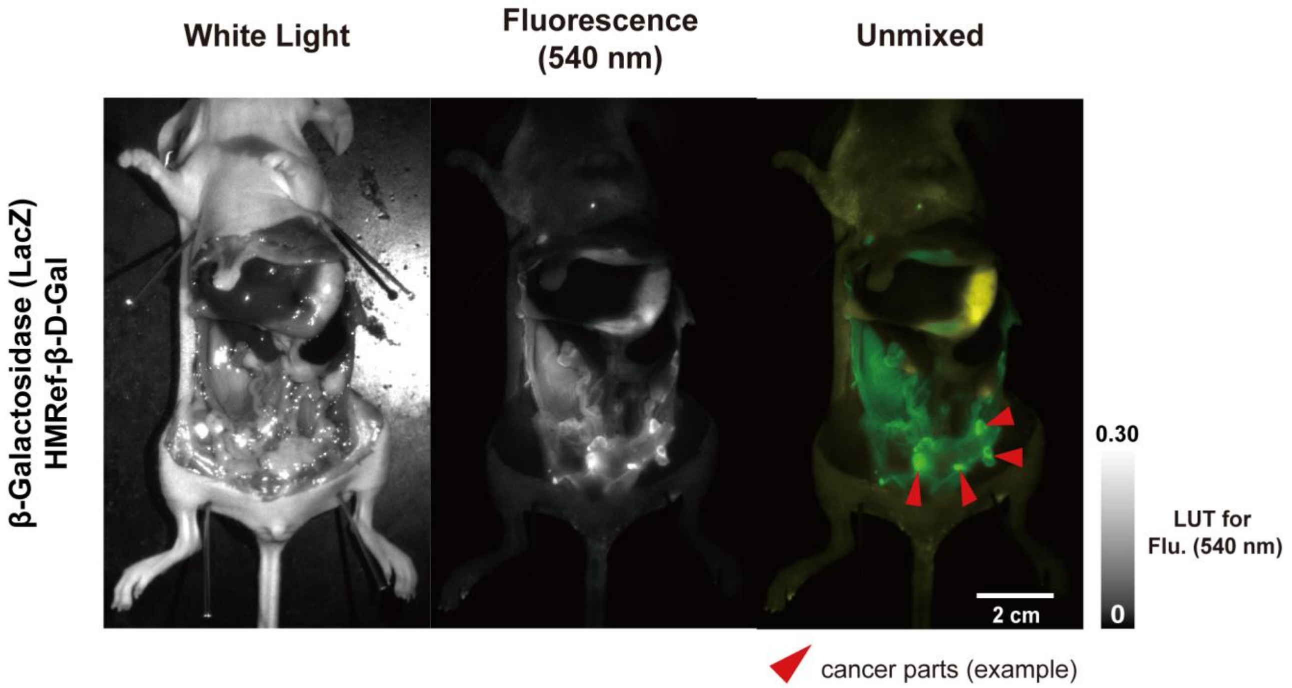


**Supplementary figure 10**. Fluorescence imaging of a mouse with intraperitoneal dissemination of SKOV-3-luc cells by using the β-galactosidase system. After the model was developed, 250 μL of 1 μM trastuzumab-β-galactosidase conjugate was i.p. injected. Three hours later, 150 μL of 1 mM HMRef-β-D-Gal was i.p. injected, and the mouse was subjected to imaging after 1 hour. The intestines were dissected out before this imaging. Fluorescence: Ex/Em = 490 nm/515 nm-longpass, exposure time = 500 ms. The stage was set at position 2, and the lighting at position 2. Scale bar = 2 cm. LUT: look up table.
